# A Trisomy 21-linked Hematopoietic Gene Variant in Microglia Confers Resilience in Human iPSC Models of Alzheimer’s Disease

**DOI:** 10.1101/2024.03.12.584646

**Authors:** Mengmeng Jin, Ziyuan Ma, Rui Dang, Haiwei Zhang, Rachael Kim, Haipeng Xue, Jesse Pascual, Steven Finkbeiner, Elizabeth Head, Ying Liu, Peng Jiang

**Author notes:** Address correspondence to: Peng Jiang, Ph.D., Associate Professor, Department of Cell Biology and Neuroscience Rutgers University New Brunswick, 604 Allison Road, Piscataway, NJ 08854.

## Abstract

**SUMMARY:** While challenging, identifying individuals displaying resilience to Alzheimer’s disease (AD) and understanding the underlying mechanism holds great promise for the development of new therapeutic interventions to effectively treat AD. Down syndrome (DS), or trisomy 21, is the most common genetic cause of AD. Interestingly, some people with DS, despite developing AD neuropathology, show resilience to cognitive decline. Furthermore, DS individuals are at an increased risk of myeloid leukemia due to somatic mutations in hematopoietic cells. Recent studies indicate that somatic mutations in hematopoietic cells may lead to resilience to neurodegeneration. Microglia, derived from hematopoietic lineages, play a central role in AD etiology. We therefore hypothesize that microglia carrying the somatic mutations associated with DS myeloid leukemia may impart resilience to AD. Using CRISPR-Cas9 gene editing, we introduce a trisomy 21-linked hotspot CSF2RB A455D mutation into human pluripotent stem cell (hPSC) lines derived from both DS and healthy individuals. Employing hPSC-based *in vitro* microglia culture and *in vivo* human microglia chimeric mouse brain models, we show that in response to pathological tau, the CSF2RB A455D mutation suppresses microglial type-1 interferon signaling, independent of trisomy 21 genetic background. This mutation reduces neuroinflammation and enhances phagocytic and autophagic functions, thereby ameliorating senescent and dystrophic phenotypes in human microglia. Moreover, the CSF2RB A455D mutation promotes the development of a unique microglia subcluster with tissue repair properties. Importantly, human microglia carrying CSF2RB A455D provide protection to neuronal function, such as neurogenesis and synaptic plasticity in chimeric mouse brains where human microglia largely repopulate the hippocampus. When co-transplanted into the same mouse brains, human microglia with CSF2RB A455D mutation phagocytize and replace human microglia carrying the wildtype CSF2RB gene following pathological tau treatment. Our findings suggest that hPSC-derived CSF2RB A455D microglia could be employed to develop effective microglial replacement therapy for AD and other age-related neurodegenerative diseases, even without the need to deplete endogenous diseased microglia prior to cell transplantation.

## Introduction

Alzheimer’s disease (AD) is the most common form of dementia, accounting for at least two-thirds of dementia cases among individuals aged 65 years and older. Characteristic neuropathological features, including beta amyloid (Aβ) plaques and hyperphosphorylated tau (p-tau)-containing neurofibrillary tangles, manifest before the onset of AD-related dementia by several years to decades. This suggests the existence of various protective mechanisms that function during this phase to stave off cognitive decline ^1–3^. Despite these protective mechanisms, the majority of individuals with AD neuropathology will eventually progress to AD dementia. Interestingly, some individuals demonstrate “resilience” to the onset of AD dementia ^1,4,5^. These individuals endure the presence of Aβ and/or p-tau while evading the gradual decline of memory functions associated with AD pathology. As such, they represent an intriguing subgroup within the human population, of significant interest for both clinical and basic research. Unveiling the mechanisms that confer resilience against AD dementia holds promise for the development of therapeutic interventions to effectively treat AD.

Remarkably, recent studies suggest a notable possibility: both the initiation of and resilience to neurodegeneration may stem from hematopoietic sources. For instance, analyzing blood DNA sequencing data from elderly individuals with and without AD, a recent study has revealed that clonal hematopoiesis, signifying a premalignant expansion of mutated hematopoietic stem cells (HSCs), is associated with protection against AD^6^. Marrow-derived myeloid cells carrying the potentially protective gene mutations likely infiltrate selective brain regions due to enhanced permeability of the blood-brain barrier under neurodegenerative conditions, exerting resilience effects to neurodegeneration ^6^. Conversely, in the context of histiocytic disorders, circulating myeloid cells with disease-causing mutations exhibit senescent phenotypes, contributing to the breakdown of the blood-brain barrier. Upon infiltrating the brain, these myeloid cells differentiate into senescent microglia-like macrophages, causing neurodegeneration^7^. Down syndrome (DS), resulting from the triplication of human chromosome 21 (Hsa21), represents the most significant genetic risk factor for AD ^8–10^. This heightened risk is primarily attributed to the overexpression of the amyloid precursor protein (APP), whose gene is situated on Hsa21, leading to early Aβ accumulation in individuals with DS in their early 30s and occasionally during childhood ^9,11^. Although individuals with DS universally develop both Aβ plaques and neurofibrillary tangles by the age of 40 years ^12–14^ and most progress to AD dementia by the age of 60 years, there exists a subset (approximately 10%) that demonstrates resilience and does not exhibit signs of dementia even at 65 years of age or above ^9,15,16^. Notably, one intriguing aspect of DS is the heightened risk of developing myeloid leukemia ^17–20^. Myeloid leukemia (ML) in DS is preceded by a preleukemic stage known as transient abnormal myelopoiesis (TAM). Approximately 30% of neonates with DS are affected by TAM ^21,22^. Moreover, 20% of DS children with TAM (approximately 6% of all DS individuals) progress to ML-DS within the first 5 years of life due to acquiring somatic mutations in HSCs ^23,24^, with an overall high cure rate (80–100%)^25^. Could the resilience to AD in the subset of DS individuals partly stem from the mutations in HSCs and their derivative myeloid cells?

Recent studies have underscored that microglial dysfunction is a central mechanism in AD etiology ^26–28^. Interestingly, the mutations found in the blood of individuals with a decreased risk of AD were also identified in the microglia-enriched fraction of their brains ^6^. While that study reported various mutations in microglia ^6^, it was not able to specifically identify a mutation that unequivocally imparts a protective mechanism against AD. A recent study highlights that a hotspot gain-of-function mutation in the myeloid cytokine receptor CSF2RB (A455D/T) plays a critical role in preleukemic and leukemic transformation in DS ^29^. *Colony stimulating factor 2 receptor subunit β* (*CSF2RB*) encodes the common β chain of the receptors for interleukin-3 (IL-3), IL-5, and granulocyte-macrophage CSF (GM-CSF). CSF2RB forms complexes with receptor-specific α chains and cytokines, triggering signaling through Janus kinase (JAK) and downstream STAT5, PI3K-AKT-mTOR, and MEK/ERK pathways to promote hematopoietic cell survival, proliferation, and differentiation ^30^. The gain-of-function CSF2RB A455D/T variant induces ligand-independent STAT5 phosphorylation, fostering cytokine-independent myeloid cell growth ^29^. In microglia, activation of CSF2RB promotes their proliferation, and GM-CSF can function as a signaling molecule stimulating innate immunity ^31^. Intriguingly, GM-CSF is upregulated in individuals with rheumatoid arthritis and associated with their protection from AD ^32^. Previous studies have also demonstrated that GM-CSF treatment yields protective effects and enhances cognitive functions in mouse models of AD ^32,33^, mouse models of DS ^34^, as well as in aged wild-type mice ^35^. A recent phase II clinical trial study has reported that GM-CSF treatment stimulates the innate immune system, improves cognition, and normalizes plasma biomarkers of neuropathology in AD patients ^36^. Taken together, based on i) the crucial functions of CSF2RB in microglia; ii) the evident beneficial effects of stimulating the innate immune system through GM-CSF treatment in both preclinical and clinical AD studies; and iii) the coincidentally similar rate (∼6% vs. ∼10%) of developing ML and acquiring resilience to AD in the DS population, we propose a hypothesis that microglia carrying the trisomy 21-associated gain-of-function mutation in CSF2RB may impart resilience to AD.

Given the limited availability of functional brain tissue from individuals with AD resilience, especially at advanced ages, and the incomplete recapitulation of human disease features in AD rodent models ^37,38^, it is challenging to validate hypothesized AD-resilience gene variants. Here, we addressed these difficulties using human pluripotent stem cell (hPSC) and CRISPR-Cas9 technologies to introduce the CSF2RB A455D mutation into DS and normal hPSCs. Through the creation of xenograft-based human-mouse brain chimeras, which have demonstrated significant relevance in modeling DS and AD^37,39–43^, we show that human microglia carrying the CSF2RB A455D mutation provide neuroprotection and resilience against pathological tau-induced alterations, as well as phagocytize and replace wild-type microglia developed in the same chimeric brain.

## Results

### The CSF2RB A455D mutation provides protection to DS human induced pluripotent stem cell (hiPSC)-derived microglia against *in vitro* pathological tau-induced cytotoxicity

Our previous study has reported the impacts of trisomy 21 on DS microglia development, as well as their senescent/dystrophic phenotypes in response to pathological tau proteins, utilizing hiPSC-based in vitro and in vivo chimeric brain models^42^. To examine the impact of the CSF2RB A455D mutation on DS microglia, we utilized CRISPR–Cas9-mediated gene editing to generate DS hiPSC lines carrying the CSF2RB A455D (DS-A455D) mutation (Figure 1A and Figure S1). Subsequently, we differentiated DS WT and DS A455D hiPSCs into primitive macrophage progenitor (PMP) cells, denoted as DS-WT and DS-A455D PMPs. As shown in Figure 1, over 95% of both WT and A455D DS hiPSC-derived PMP expressed CD235 (a yolk sac primitive hematopoietic progenitor marker) and CD43 (a hematopoietic progenitor-like cell marker). Both DS-WT and DS-A455D PMPs exhibited high proliferative activity, evident from the majority of cells expressing Ki67 (Figure 1B, 1C). To investigate potential transcriptomic changes induced by the A455D mutation in DS PMPs, we conducted RNA-seq analysis on DS-WT and DS-A455D PMPs. As anticipated, the A455D mutation was absent in DS-WT PMPs but present in more than 95% of DS-A455D PMPs (Figure 1D). Correlation analyses revealed robust similarities between DS-WT and DS-A455D PMPs at the whole transcriptome level (Spearman’s ρ = 0.9237, Figure 1E), suggesting that A455D does not significantly alter PMP identity. Additionally, no significant differences in gene expression were observed between DS-WT and DS-A455D PMP (Figure 1F). The A455D variant is known to induce an active conformation for downstream JAK signaling, leading to the constitutive phosphorylation of STAT5 ^29^. Consistently, DS-A455D PMPs exhibited a significantly higher level of phosphorylated STAT5 than DS-WT PMPs, as evidenced by flow cytometry targeting pSTAT5 under a cytokine-free condition (Figure 1G, 1H). These findings indicate that the CSF2RB A455D mutation induces the phosphorylation of STAT5 but does not significantly alter the transcriptome of DS PMPs.

**Fig 1.**
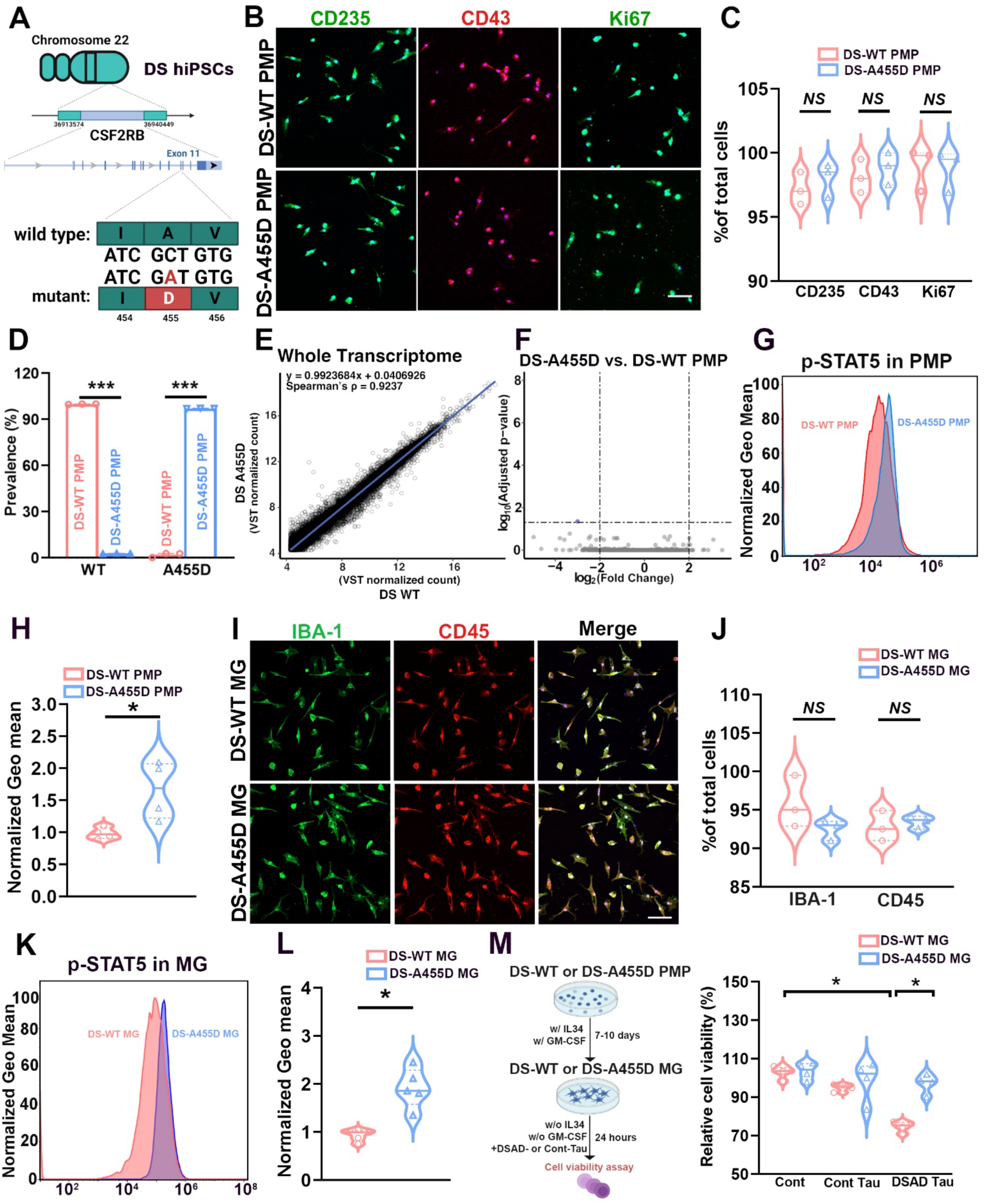
Generation and characterization of DS-WT and DS-A455D hiPSC-derived PMPs, and the tau-induced cytotoxicity on DS-WT and DS-A455D MG. (A) Schematic representation of the generation and characterization of DS-A455D hiPSCs. (B) Representative images of CD235^+^, CD43^+^, and Ki67^+^ cells in DS-WT and DS-A4555D PMPs. Scale bars: 40 μm. (C) Quantification of CD235^+^, CD43^+^, and Ki67^+^ PMPs derived from the DS-WT and DS-A455D hiPSC lines (n = 3, each experiment was repeated three times). Student’s t-test, *NS*, not significant. Data are presented as mean ± SEM. (D) The A455D prevalence in DS-WT and DS-A455D PMP (n = 3, each experiment was repeated three times). Student’s t-test, \*\*\**P* < 0.001. Data are presented as mean ± SEM. (E) Linear regression and Spearman’s correlation tests of bulk RNA-seq data between DS-WT and DS-A455D PMP at the whole transcriptome level. (F) A volcano plot comparing genes expressed in DS-WT and DS-A455D PMP. (G) Flow cytometry analysis of p-STAT5 level in DS-WT and DS-A455D PMP. (H) Quantification of p-STAT5 level in DS-WT and DS-A455D PMP (n = 4). Student’s t-test, \**P* < 0.05. Data are presented as mean ± SEM. (I) Representative images of IBA-1^+^ and CD45^+^ in DS-WT and DS-A4555D MG. Scale bars: 40 μm. (J) Quantification of IBA-1^+^ and CD45^+^ MG derived from the DS-WT and DS-A455D hiPSC lines (n = 3, each experiment was repeated three times). Student’s t-test, *NS*, not significant. Data are presented as mean ± SEM. (K) Flow cytometry analysis of p-STAT5 level in DS-WT and DS-A455D MG. (L) Quantification of p-STAT5 level in DS-WT and DS-A455D MG (n = 5). Student’s t-test, \**P* < 0.05. Data are presented as mean ± SEM. (M) Schematic representation of *in vitro* tau-induced cytotoxicity on DS-WT and DS-A455D MG created with BioRender.com. Quantitative analysis of cell viability after Tau treatment (n = 4). Two-way ANOVA test, \**P* < 0.05. Data are presented as mean ± SEM.

To determine the functionality of CSF2RB A455D in microglia, we next differentiated these PMPs into microglia *in vitro*, referred to as DS-WT and DS-A455D microglia (MG). The differentiated cells universally expressed microglia/macrophage markers CD45 and IBA-1, with no discernible differences between DS-WT and DS-A455D MG (Figure 1I, 1J). Consistent with the PMP stage, flow cytometry analysis revealed elevated levels of STAT5 phosphorylation in DS-A455D MG compared to DS-WT MG, confirming the constitutive activation of pSTAT5 in microglia (Figure 1K, 1L). Previous *in vitro* studies have demonstrated that soluble hyperphosphorylated tau (p-Tau) extracted from human AD brain tissue induces microglial cell death ^44^. We then evaluated the impact of the S1 soluble fractions containing only p-tau, but not Aβ derived from DSAD human brain tissue (DSAD-Tau) ^42,44^ on DS-WT and DS-A455D MG. After subjecting microglia to a 24-hour treatment, a cell viability test revealed that DSAD-Tau exerted a toxic effect on DS-WT MG, whereas DS-A455D MG were protected from DSAD-Tau-induced cytotoxicity (Figure 1M). The cell viability of DS-WT and DS-A455D MG remained unaffected by the soluble S1 fraction isolated from age-and sex-matched control brain tissue (Cont-Tau). Hence, these findings establish that the CSF2RB A455D mutation provides protection to DS microglia against p-tau-induced cytotoxicity *in vitro*.

### DS-A455D MG develop a distinct protective microglial subpopulation *in vivo* in response to pathological tau

Our previous study^42^ demonstrated that human microglia chimeric mouse brain model faithfully recapitulated DS microglia phenotypes reported in mouse models of DS ^45^ as well as dystrophic phenotypes observed in DS postmortem human brain tissue ^46,47^. To better mature the DS PMPs into microglia and examine the effects of CSF2RB A455D *in vivo*, we transplanted DS-A455D and DS-WT PMPs into the hippocampus of postnatal day 0 (P0) Rag2^−/−^ IL2rγ^−/−^ hCSF1^KI^ immunodeficient mice, establishing human microglia mouse chimeras ^42,48,49^. Eight weeks post-transplantation, the vast majority of hN^+^ donor-derived cells (>90%) in both DS-WT and DS-A455D groups were hTMEM119^+^ (Figure S2A, S2C), indicating that DS-A455D and DS-WT PMPs similarly give rise to microglia *in vivo*. In the human microglia chimeric mouse brains, both DS-WT and DS-A455D MG remained proliferative at 2 months post-transplantation, and no difference in Ki67^+^ cells among total hN^+^ donor-derived cells was observed (Figure S2B, S2D). We then analyzed the level of pSTAT5 by flow cytometry targeting human specific CD45^+^ (hCD45^+^) microglia in these chimeras. Consistently, DS-A455D chimeras exhibited higher levels of pSTAT5 expression than DS-WT chimeras (Figure S2E, S2F), aligning with the constitutive activation of CSF2RB and pSTAT5 signaling due to the CSF2RB A455D mutation ^29^. To investigate the responses of DS-A455D microglia to pathological tau proteins *in vivo*, these chimeric mice at the age of 2 months received injections of either Cont-Tau or DSAD-Tau. At 2 months following tau injection (4 months of age), hTMEM119^+^ DS-A455D and DS-WT MG were widely distributed throughout the hippocampus (Figure 2A). Some hN^+^/hTMEM119^-^human cells were observed in the lateral ventricles, likely representing brain border-associated macrophages differentiated from engrafted PMPs, as shown in previous studies ^40,48^. The hTMEM119^+^ DS-A455D MG responded to injected p-tau, displaying a significantly greater amount of engulfed AT8^+^ p-tau in the DSAD-Tau group (DS-A455D-DSAD-Tau group) than in the Cont-Tau group (DS-A455D-Cont-Tau group) (Fig 2B, 2C). Additionally, these DS-A455D MG exhibited an enhanced proliferation in response to injected p-tau, as indicated by a higher percentage of Ki67^+^/hN^+^ cells in the DS-A455D-DSAD-Tau group compared to the DS-A455D-Cont-Tau group (Figure S2G, S2H).

**Fig 2.**
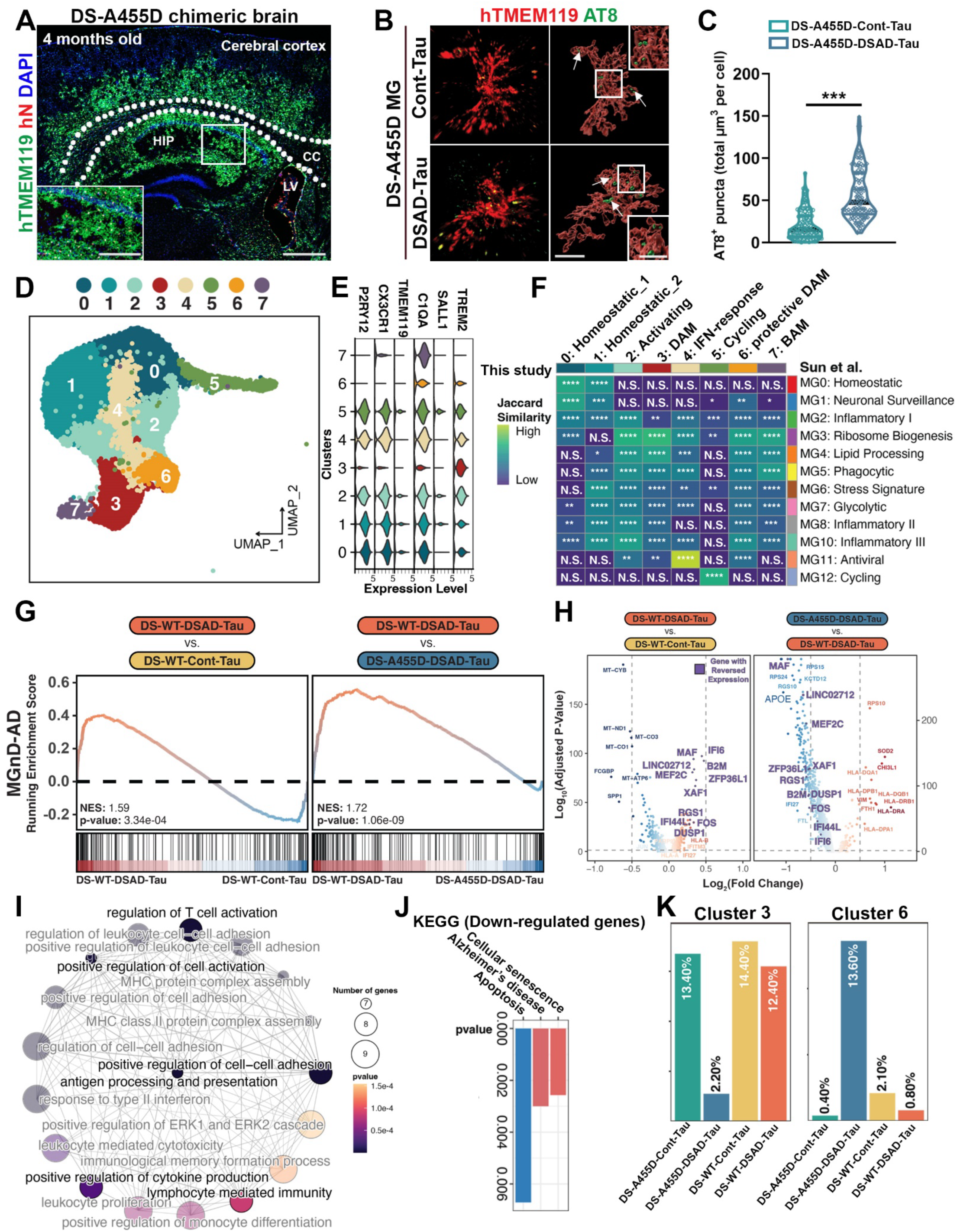
DS-A455D MG give rise to a distinct microglial subpopulation in response to pathological tau in chimeric brains. (A) Representative images from sagittal brain sections showing the distribution of transplanted DS-A455D MG. Scale bar: 500 and 200 μm in the original and enlarged images, respectively. (B) Representative raw fluorescence super-resolution and 3D surface rendered images showing colocalization of hTMEM119^+^ and AT8^+^ staining in 4-month-old chimeric mice receiving injection of Cont or DSAD Tau at the age of 8 weeks. Scale bars: 5 μm and 3 μm in the original and enlarged images, respectively. (C) Quantification of AT8^+^ p-tau in DS-A455D microglia following Cont and DSAD Tau injection (n = 121-132 from 3 mice per group). Student’s t-test, \*\*\**P* < 0.001. Data are presented as mean ± SEM. (D) A UMAP plot showing microglia subclusters (clusters 0-7) from Cont-Tau and DSAD-Tau groups. (E) A violin plot showing the expression of human microglial marker genes from each subcluster. (F) Heatmap to show the similarity of clusters between Sun et al. (human) and this study. The Jaccard score represents the percentage of pairwise overlapping genes. Statistical significances of the overlappings are obtained from contingency tables using Fisher’s exact test. (G) GSEA plots showing enrichment of MGnD-AD genes in DS-WT-DSAD-Tau and DS-WT-Cont-Tau as well as DS-WT-DSAD-Tau and DS-A455D-DSAD-Tau groups (NES: normalized enrichment score). (H) Volcano plots depict downregulated genes (blue) and upregulated genes (red) for both DS-WT-DSAD-Tau vs. DS-WT-Cont-Tau MG and DS-A455D-DSAD-Tau vs. DS-WT-DSAD-Tau MG comparisons. In the latter group, reversed gene names are color-coded in purple. (I) Enrichment map showing GO-BP enrichment analysis of the upregulated DEGs in DS-A455D-DSAD-Tau vs. DS-WT-DSAD-Tau MG group. (J) KEGG enrichment analysis of the downregulated DEGs in DS-A455D-DSAD-Tau vs. DS-WT-DSAD-Tau MG group. (K) Bar plots showing cell proportions of clusters 3 and 6 in all experimental groups.

To characterize the responses of DS-A455D MG to pathological tau *in vivo* at the molecular level, we performed single-cell RNA sequencing (scRNA-seq) using DS-A455D-DSAD-Tau and DS-A455D-Cont-Tau chimeric mice (Figure S2I). The data were integrated with our prior scRNA-seq analysis of DS-WT microglia chimeras exposed to either Cont-Tau (DS-WT-Cont-Tau) or DSAD-Tau (DS-WT-DSAD-Tau) ^42^. After excluding mouse cells and implementing quality control measures (method, Figure S2J), we analyzed 25,631 human microglia. Shared nearest neighbor (SNN) clustering revealed 10 clusters (Figure S2K), including 8 microglial clusters (clusters 0-7, characterized by *P2RY12, CX3CR1, TMEM119, C1QA, SALL1*, and *TREM2*, Figure 2D) and 2 unidentified clusters (cluster 8, characterized by *COL3A1, COL1A1,* and *COL1A2*, and cluster 9, by *TTR, CLU,* and *IGFBP7*, Figure S3A). Owing to their absence of canonical microglial markers (Figure S3B), low similarity with other microglial clusters (Figure S3C), and representation of a small portion of total human cells (∼1.5%), clusters 8 and 9 were excluded from further analysis. The remaining 8 microglial clusters, encompassing 25,235 cells, were further classified into distinct states based on their transcriptomic profiles. Figure 2E depicts a violin plot illustrating the expression of human microglial marker genes in each cluster. Clusters 0 and 1 represented homeostatic microglia, indicated by abundant expression of *P2RY12, CX3CR1,* and *TMEM119*. Cluster 2 signified a microglial population transitioning from a homeostatic to an activated state. Clusters 3 and 6 were identified as disease-associated microglia (DAM), characterized by enriched expression of DAM-related genes, ferritin gene expression, and phagocytic signatures ^50–52^ (Figure S3D). Cluster 4 was distinguished as an interferon (IFN) response-related microglia cluster due to strong expression of IFN-related genes (e.g., *IFIT1, ISG15,* and *IFIT3*). Cluster 5, with high expression of cell-cycle related genes (e.g., *MKI67, CENPF,* and *TOP2A*) and a high cell cycle score (Figure S3E), was termed as a cycling microglia cluster. Cluster 7 was identified as a population of brain border-associated macrophages, marked *by MRC1, CD163,* and *LYVE1*^53^. This cluster likely corresponds to the hN^+^/hTMEM119^-^human cells observed in lateral ventricles (Figure 2A).

We next conducted a Jaccard similarity analysis to compare our data with microglia single-nucleus RNA-seq (snRNA-seq) data from postmortem AD human brain tissue recently reported by Sun et al. ^54^ (Figure 2F). Our homeostatic microglial clusters 0 and 1 exhibited high Jaccard similarity with Sun et al.’s homeostatic cluster MG0. The IFN microglial cluster 4 closely corresponded to antiviral MG11 described by Sun et al.’s, and our cycling microglial cluster 5 shared a gene signature with Sun et al.’s cycling cluster MG12. Our two DAM clusters also showed significant overlap in marker genes related to ribosome biogenesis, lipid processing, phagocytosis, stress response, glycolysis, and inflammatory processes, aligning with DAM microglia states in AD patients’ brains ^54^ (Figure 2F). These findings suggest that DS microglia, when exposed to pathological tau derived from human patients in chimeric mouse brains, exhibit overlapping transcriptomic signatures and heterogeneity to microglia in AD patients in the general population. We then performed a gene set enrichment analysis (GSEA) to capture the expression profiles of AD-associated neurodegenerative microglia (MGnD) ^50^. We observed the majority of lower enrichment of MGnD-AD signature genes were enriched in the DS-WT-DSAD-Tau group instead of the DS-A455D-DSAD-Tau group (Figure 2G). Differentially expressed gene (DEG) analysis showed that genes upregulated in the DS-WT-DSAD-Tau group, in contrast to DS-WT-Cont-Tau group, were reversed in the DS-A455D-DSAD-Tau group (Figure 2H). Furthermore, gene ontology (GO) enrichment analysis showed enrichment in biological process terms related to the regulation of T cell activation, antigen processing and presenting, and regulation of cytokine production in the DS-A455D-DSAD-Tau group (Figure 2I). Notably, by performing Kyoto Encyclopedia of Genes and Genomes (KEGG) enrichment analyses, we found that many significantly enriched DEGs downregulated in the DS-A455D-DSAD-Tau group were associated with cell senescence pathways (e.g., *B2M, ZFP36L1*, and *XAF1*), Alzheimer’s Disease (e.g., *APOE* and *TREM2*), and apoptosis pathways (Figure 2J). Interestingly, within the two DAM clusters in our dataset, cluster proportion analysis revealed distinct sample distributions among the four groups. DAM cluster 3 exhibited a significantly reduced proportion in the DS-A455D-DSAD-Tau group compared to the other three groups (Figure 2K). In contrast, DAM cluster 6 displayed the highest proportion in the DS-A455D-DSAD-Tau group (Figure 2K). Importantly, the cells identified as cluster 6 cells predominantly originated from the DS-A455D-DSAD-Tau group, suggesting that cluster 6 was likely uniquely associated with the CSF2RB A455D mutation.

Although both clusters 3 and 6 significantly displayed DAM signatures, they possessed distinct DAM marker gene expression profiles (Figure 3A). To further investigate these differences, we conducted pseudo-time trajectory inference for the DS-A455D-DSAD-Tau group. This analysis predicted five trajectories originating from homeostatic microglia cluster 0 (Figure S4A), with two trajectories of particular interest: lineage 1 leading to DAM cluster 6 and lineage 3 leading to DAM cluster 3 (Figure 3B). For a detailed analysis of gene expression changes in lineage 1 and 3, we calculated dynamically changed DEGs across the trajectories, and applied K-Nearest Neighbors (KNN) clustering to discern expression patterns across 101 and 149 variably expressed genes (Figure 3C, 3D, Supplementary Table S6-7). GO enrichment analysis on these KNN clusters revealed intriguing findings. KNN cluster C5 showed enrichment in biological process terms related to plasma membrane organization, repair, tissue remodeling, wound healing, and lipid transport, suggesting a protective microglial state of DAM cluster 6 (Figure 3C, Supplementary Table S8). In contrast, KNN cluster C3 showed enrichment in biological process terms related to lipid-droplet-accumulation and antigen processing and presenting, which represented a dysfunctional microglia state consistent with a previous study ^55^ (Figure 3D, Supplementary Table S9). Next, we analyzed DEGs between cluster 6 and cluster 3 in DS-WT-DSAD-Tau and DS-A455D-DSAD-Tau groups. The volcano plot showed top upregulated genes (e.g., *CHI3L1, VIM*, and *SOD2*) that were related to tissue remodeling, and wound healing (Figure 3E, Supplementary Table S10). For a detailed analysis of DEGs in clusters 3 and 6, we further plotted the expression changes of on some top DEGs across pseudotime (Figure 3F, S4B). Many genes linked to tissue remodeling and repair exhibited more pronounced expression in cluster 6 than in cluster 3. For instance, chitinase-3–like protein 1 (CHI3L1) is a master regulator for a wide range of injury and repair responses, including the innate immunity pathways in microglia ^56,57^. Pseudo-time trajectory inference on *CHI3L1* in lineage 1 and 3 unveiled a more significant expression in cluster 6 compared to cluster 3. Another example is superoxide dismutase 2 (*SOD2*), which is also expressed at a higher level in cluster 6 than in cluster 3. SOD2, a major antioxidative enzyme, has been implicated in mitigating inflammatory reactions, owing to its suppressive effects on NF-κB activity and the production of proinflammatory cytokines ^58^. Sequestosome 1 (SQSTM1 or p62), henceforth referred to as p62, plays a key role in directing ubiquitinated proteins to the proteasome or the growing autophagosome for subsequent protein degradation ^59^. P62 is critically involved in tau degradation and the frontal cortex of AD patients and transgenic AD mouse models exhibited low p62 expression ^60^. Our scRNA-seq analysis revealed a high expression of *p62* in cluster 6 compared to cluster 3. Overall, these findings underscore that the CSF2RB A455D mutation actively contributes to protecting DS microglia from p-tau *in vivo*, with DAM cluster 6 identified as a protective cluster associated with the CSF2RB A455D mutation.

**Fig 3.**
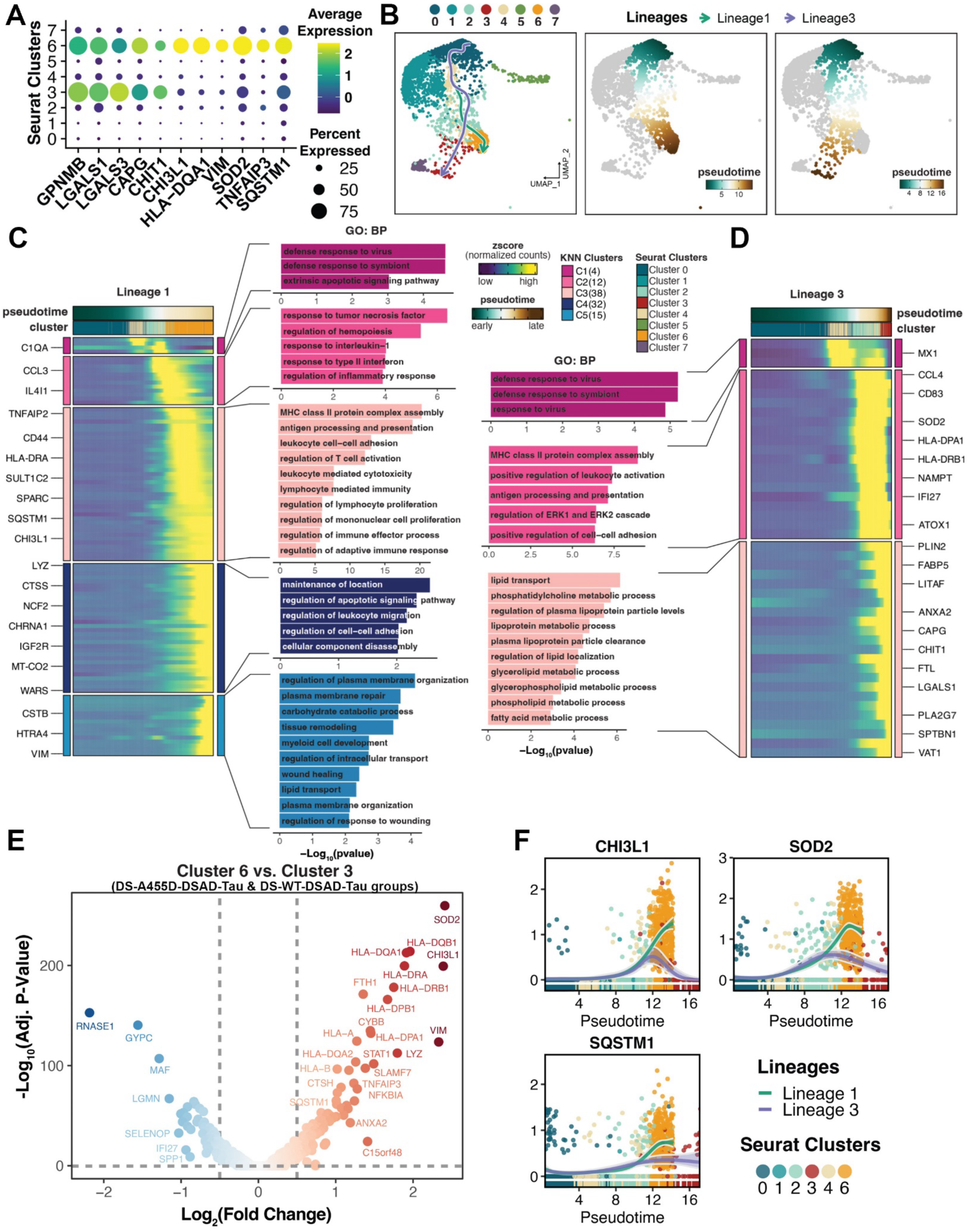
DAM clusters identified from DS-A455D microglial chimeric brains receiving injection of Cont or DSAD tau. (A) A dot plot showing marker gene expression profiles of DAM cluster 3 and 6. (B) UMAP representation 2 of the 5 inferred trajectories of DS-A455D MG in response to DSAD-Tau. Cells are colored by microglial subclusters in the left panel and by pseudotime in the middle and right panels. (C, D) Dynamic heatmap representation of DEG expression profiles of lineage 1 (C) and lineage 3 (D) in DS-A455D-DSAD-Tau group across pseudotime. And GO-BP enrichment analyses of each KNN cluster of lineage 1 and 3. (E) A volcano plot showing the downregulated (blue) and upregulated (red) DEGs in DSAD-Tau-treated groups by comparing cluster 6 vs. cluster 3. (F) Dynamic plots showing the selected gene expression profiles across inferred pseudotime in lineage 1 and lineage 3. Dots in the plots are cells colored by microglial subclusters.

### The CSF2RB A455D mutation protects DS microglia against senescence

Our previous study revealed that exposure to pathological tau leads to an upregulation of type-I interferon (IFN-I) signaling in DS-WT MG, contributing to accelerated senescence of DS microglia ^42^. In this study, GSEA analysis indicated significantly lower senescence gene enrichment in the DS-A455D-DSAD-Tau group compared to the DS-WT-DSAD-Tau group (Figure 4A). Specifically, DAM clusters 3 and 6 in the DS-A455D-DSAD-Tau group exhibited significant lower senescence scores than those in the DS-WT-DSAD-Tau group (Figure 4B). By comparing the DEGs between the DS-A455D-DSAD-Tau group and the DS-WT-DSAD-Tau group, we found 41 genes are type-1 IFN-related. Notably, the majority of these genes (38 genes, 92.68%) were down-regulated in the DS-A455D-DSAD-Tau group (Figure 4C). Double-staining for IBA-1 and hN revealed that hN^+^/IBA-1^+^ DS-A455D microglia in the DSAD-Tau group displayed normal processes and less fragmentation compared to the dystrophic morphology of DS-WT microglia in the DSAD-Tau group (Figure 4D). Quantitative analysis showed increased microglia volume, prolonged process length, decreased soma size, and a reduced ratio between soma size and process length in the DS-A455D-DSAD-Tau group compared to the DS-WT-DSAD-Tau group (Figure 4F-I). Subsequent staining of chimeric mouse brain tissue with hCD45 and ferritin, a marker of senescent microglia, demonstrated fewer ferritin^+^/hCD45^+^ microglia in the DS-A455D-DSAD-Tau group than in the DS-WT-DSAD-Tau group (Figure 4E, 4J). These experimental data further support that the CSF2RB A455D mutation protects DS microglia from undergoing senescence.

**Fig 4.**
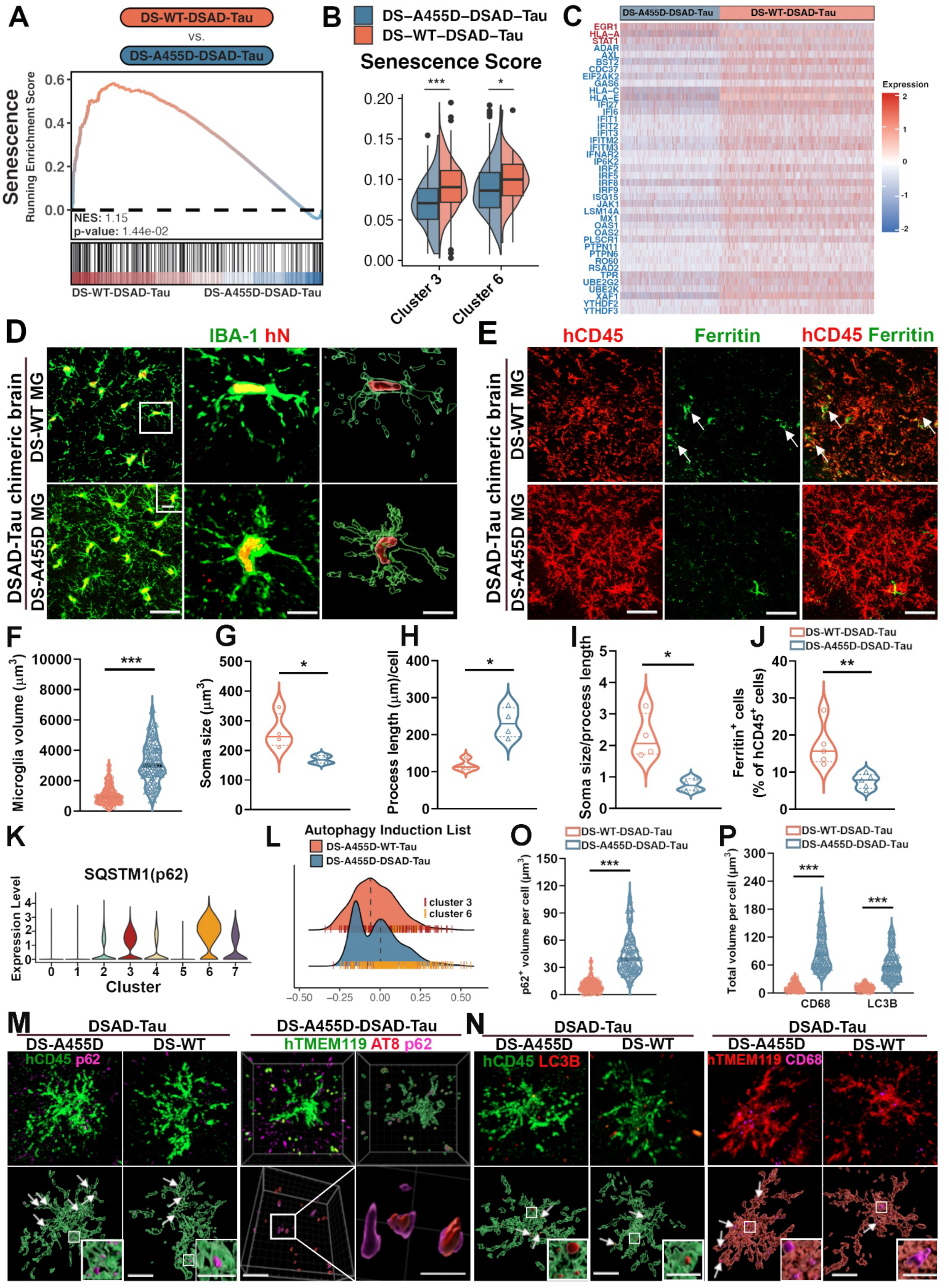
CSF2RB A455D mutation protects DS microglia against senescence. (A) GSEA plot showing enrichment of senescence genes in DS-WT-DSAD-Tau and DS-A455D-DSAD-Tau groups. (B) Violin plots showing the senescence score of clusters 3 and 6 from DS-WT-DSAD-Tau and DS-A455D-DSAD-Tau groups. Wilcoxon test, \**P* < 0.05, \*\*\**P* < 0.001. (C) Heatmap representation of IFN-related genes in the DS-A455D-DSAD-Tau and DS-WT-DSAD-Tau groups. Red: down-regulated genes in DS-WT-DSAD-Tau group; Blue: down-regulated genes in DS-A455D-DSAD-Tau group. (D) Representative images of IBA^+^/hN^+^ human microglia in DS-WT-DSAD-Tau and DS-A455D-DSAD-Tau groups. Scale bars: 20 μm and 5 μm. (E) Representative images showing colocalization of hCD45^+^ and Ferritin^+^ staining in DS-WT-DSAD-Tau and DS-A455D-DSAD-Tau groups. Arrows indicate Ferritin^+^ and/or hCD45^+^ staining. Scale bar: 20 μm. (F) Quantification of the microglia volumes in DS-WT-DSAD-Tau and DS-A455D-DSAD-Tau groups (n = 119-122 from 3 mice per group). Student’s t-test, \*\*\**P* < 0.001. Data are presented as mean ± SEM. (G-I) Quantification of the process length, soma size, soma size/process length (n = 4). Student’s t-test, \**P* < 0.05. Data are presented as mean ± SEM. (J) Quantification of the percentage of Ferritin in hCD45^+^ cells (n = 5). Student’s t-test, \*\**P* < 0.01. Data are presented as mean ± SEM. (K) A violin plot showing the expression of SQSTM1 from each subcluster. (L) A ridgeline plot showing autophagy induction genes expression of clusters 3 and 6 in DS-A455D-DSAD-Tau and DS-WT-DSAD-Tau groups. (M) Representative images of hCD45 p62 staining in 4-months-old DS-A455D-DSAD-Tau and DS-WT-DSAD-Tau chimeras and hTMEM119, AT8, and p62 staining in 4-months-old DS-A455D-DSAD-Tau group. Arrows indicate p62^+^ puncta. Scale bars:5 μm and 2 μm in the original and enlarged images, respectively. (N) Representative images of hCD45 LC3B and hTMEM119 CD68 staining in 4-months-old DS-A455D-DSAD-Tau and DS-WT-DSAD-Tau chimeras. Arrows indicate LC3B^+^ and CD68^+^ puncta. Scale bars:5 μm and 2 μm in the original and enlarged images, respectively. (O) Quantification of p62^+^ puncta in hCD45+microglia (n = 130-131 microglia from 3-4 mice per group). Student’s t-test, \*\*\**P* < 0.001. Data are presented as mean ± SEM. (P) Quantification of LC3B^+^ and CD68^+^ puncta in hCD45+ microglia (n = 131 microglia from 3 mice per group). Student’s t-test, \*\*\**P* < 0.001. Data are presented as mean ± SEM.

Recent research indicates that upregulated autophagy prevents microglial senescence ^61^. To explore the mechanism behind the protective effects of the CSF2RB A455D mutation in microglia, we analyzed scRNA-seq of microglia in DS-WT-DSAD-Tau and DS-A455D-DSAD-Tau groups. Very interestingly, the autophagy adaptor protein p62 was expressed in both DAM groups and highly expressed in DAM clusters 3 and 6 (Figure 4K), consistent with prior reports linking microglial autophagy activation with DAM ^61^. Gene set scoring of DAM clusters 3 and 6 revealed a higher score of autophagy induction genes in the DS-A455D-DSAD-Tau group (dominated by cells originated from cluster 6) compared to the DS-WT-DSAD-Tau (dominated by cells originated from cluster 3, Figure 4L). Next, we assessed p62 expression in hCD45^+^ microglia in DS-WT-DSAD-Tau and DS-A455D-DSAD-Tau groups using double staining for hCD45 and p62. p62^+^ puncta were observed in both groups, but 3D-reconstruction imaging showing significantly more p62^+^ puncta in DS-A455D-DSAD-Tau microglia than in DS-WT-DSAD-Tau microglia (Figure 4M, 4O). Additionally, colocalization of AT8 with p62 puncta in DS-A455D-DSAD-Tau microglia was observed (Figure 4M). Furthermore, we stained hCD45 and LC3B (a phagophore and autophagosomal membrane protein, also known as MAP1LC3B-II), hTMEM119, and CD68, a marker for phagolysosomes. As shown in Figure 4N, LC3B^+^ puncta were present inside the hCD45^+^ microglia in DS-WT-DSAD-Tau and DS-A455D-DSAD-Tau groups. Microglia in the DS-A455D-DSAD-Tau group also exhibited a higher volume of LC3B^+^ autophagosomes and a higher volume of CD68^+^ phagolysosomes (Figure 4N, 4P). These results suggest that DS-A455D MG, in contrast to DS-WT MG, exhibit improved uptake of AT8^+^ DSAD-Tau proteins and subsequent degradation through the autophagy-lysosome pathway, with the activation of autophagy further aiding in preventing microglia senescence.

### DS-A455D MG safeguards neuronal functions against pathological tau

Our previous study uncovered a significant increase in *β-2 microglobulin* (*B2M)* expression in DS-WT MG following exposure to pathological tau ^42^. Interestingly, in this study, scRNA-seq analysis revealed *B2M* downregulation in DS-A455D-DSAD-Tau, compared to the DS-WT-DSAD-Tau group (Figure 5A). Flow cytometry targeting hCD45^+^ microglia cells from chimeric brains validated this finding, confirming lower B2M expression in DS-A455D MG than DS-WT MG in response to DSAD-Tau (Figure 5B, 5C). Microglia, as the resident immune cells in CNS, are implicated in neuroinflammation. Here, we observed that DS-A455D MG exhibited a significantly mitigated neuroinflammatory response and reduced expression of inflammation-related genes (e.g., *CTSC, GRN,* and *TNFRSF1B*) compared to the DS-WT MG in response to DSAD-Tau (Figure 5D, 5E).

**Fig 5.**
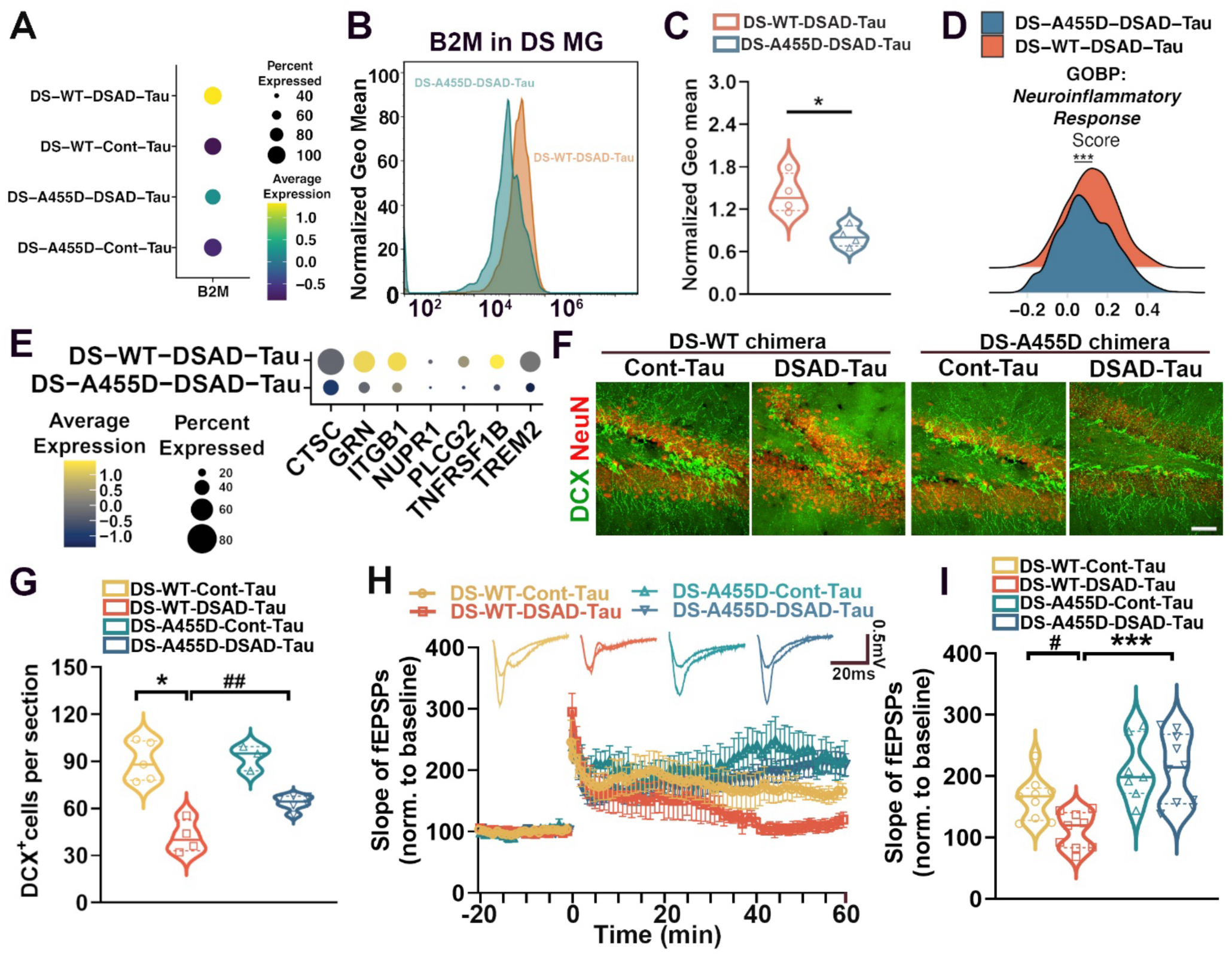
DS-A455D MG protects neuronal functions in response to pathological tau. (A) A Dot plot representing the expression profile of the B2M gene. (B) Flow cytometry analysis of B2M expression in DS-WT-DSAD-Tau and DS-A455D-DSAD-Tau groups. (C) Quantification of B2M expression in DS-WT-DSAD-Tau and DS-A455D-DSAD-Tau groups (n = 4). Student’s t-test, \**P* < 0.05. Data are presented as mean ± SEM (Geo mean: Geometric mean). (D) Ridgeline plot comparing the neuroinflammatory response gene score in DS-WT-DSAD-Tau and DS-A455D-DSAD-Tau groups. (E) Dot plot comparing selected neuroinflammatory gene expression profiles between DS-WT-DSAD-Tau and DS-A455D-DSAD-Tau group. (F) Representative images of DCX and NeuN staining in 4-months-old chimeras. Scale bars: 20 μm. (G) Quantification of DCX^+^ cells (n = 5 mice per group). Two-way ANOVA test, \**P* < 0.05, and ^##^*P* < 0.01. Data are presented as mean ± SEM. (H) Representative traces of baseline and last 10 min fEPSP after 3X 100 Hz LTP induction. Quantification of LTP after LTP induction in 4 months old chimeras (n = 7-10 slices from 3-4 mice per group). (I) Quantification of the last 10 min of fEPSP slope after LTP induction in 4 months old chimeras (n = 7-10 slices from 3-4 mice per group). Two-way ANOVA test, ^#^*P* < 0.05 and \*\*\**P* < 0.001. Data are presented as mean ± SEM.

B2M, a constituent of MHC class I molecules, has been identified as a secreted pro-aging factor primarily expressed by microglia in the brain that impairs synaptic functions, neurogenesis, and cognitive function ^62–64^. Furthermore, neuroinflammation is also known to impact neurogenesis ^65^. The findings of downregulated B2M and reduced expression of inflammation-related genes in DS-A455D MG prompted us to experimentally assess whether DS-A455D MG protect neuronal functions against pathological tau. Immunostaining analysis of hippocampal neurogenesis in adult chimeric mouse brains revealed a significantly higher number of doublecortin (DCX)-positive newborn neurons in DS-A455D-DSAD-Tau group compared to DS-WT-DSAD-Tau group (Figure 5F, 5G). Subsequently, we evaluated long-term potentiation (LTP) in the hippocampus, a cellular mechanism underlying learning and memory, in 4-month-old chimeras. LTP was induced with 3 trains of 100 Hz high-frequency stimulation (HFS). Figure 5H illustrated that DS-WT-DSAD-Tau showed impaired LTP, with reduced field excitatory postsynaptic potential (fEPSP) slope compared to the other groups (Figure 5H, 5I), whereas DS-A455D-Cont-Tau and DS-A455D-DSAD-Tau mice showed fEPSP slope comparable to DS-WT-Cont-Tau mice. As shown in Figure 5I, unlike DS-WT-DSAD-Tau mice that had impaired LTP, DS-A455D-DSAD-Tau mice exhibited LTP similar to DS-WT-Cont-Tau mice where fEPSP slope persisted for 60 min. Immunohistological staining using slices post-LTP recordings consistently displayed a wide distribution of xenografted, hTMEM119+ human microglia (Figure S5). Collectively, these results affirm that the CSF2RB A455D mutation in microglia protects neuronal functions against pathological tau.

### The resilience effects conferred by the CSF2RB A455D mutation are independent of trisomy 21

To investigate whether the protective effects of the CSF2RB A455D mutation depend on the trisomy 21 genetic background, we created two control hPSC lines harboring the CSF2RB A455D mutation – a control hiPSC line (Cont-A455D) and a hESC line (CAGG-A455D) – utilizing CRISPR–Cas-9-mediated gene editing. Subsequently, we derived PMPs from both Cont-WT and Cont-A455D iPSCs, as well as CAGG-WT and CAGG-A455D hESCs. The majority of WT and A455D hiPSC and hESC-derived PMPs expressed CD235 and CD43, demonstrating their phenotypic similarity (Figure S6A). Moreover, these PMPs exhibited high proliferation, as indicated by Ki67 expression (Figure S6A). RNA-seq analysis of the PMPs showed that more than 95% of A455D PMPs harbored the A455D variant (Figure S6B). There was a strong correlation across the entire transcriptome between all WT and A455D PMPs (Spearman’s ρ = 0.9344, Figure S6C, S6D), suggesting that A455D does not significantly alter the overall gene expression of PMPs. Flow cytometry confirmed elevated pSTAT5 activation in Cont-A455D PMPs compared Cont-WT PMPs in cytokine-free culture conditions (Figure S6E, S6F). After *in vitro* differentiation for 7-10 days, both WT and A455D microglial cells expressed microglia/macrophage markers CD45 and IBA-1, with no statistically significant differences (Figure S6G). Flow cytometry targeting pSTAT5 in cytokine-free conditions also showed significantly higher levels of STAT5 phosphorylation in A455D MG compared to WT MG, confirming pSTAT5 activation in microglia (Figure S6H, 6I). Subsequently, we exposed these microglia to soluble p-tau for 24 hours. The cell viability test demonstrated that DSAD-Tau exerted a toxic effect on WT microglia but did not significantly affect the viability of A455D microglia cells (Figure S6J).

We then implanted Cont-A455D and Cont-WT PMPs into the hippocampus of P0 immunodeficient mice to create chimeric mice. At eight weeks post-transplantation, the majority of hN^+^ donor-derived cells (>90%) in both Cont-WT and Cont-A455D groups were hTMEM119^+^ and remained proliferative (Figure S6M). Quantitative analysis revealed no difference in hTMEM119^+^ and Ki67^+^ cells among the total hN^+^ donor-derived cells (Figure S6N, S6O). Next, we examined the expression level of pSTAT5 in these chimeras. Consistently, Cont-A455D microglia chimeras exhibited a higher level of pSTAT5 expression than Cont-WT chimeras (Figure S6K, 6L). To further explore how Cont-A455D and Cont-WT MG respond to pathological tau proteins *in vivo*, these chimeric mice received injections of Cont-Tau or DSAD-Tau at the age of 2 months. At 4 months after tau injection (6 months of age), the hTMEM119^+^ Cont-A455D MG displayed many more engulfed AT8^+^ p-tau in the DSAD-Tau group than in the Cont-Tau group (Figure 6A, 6B). Additionally, the Cont-A455D MG in the Cont-A455D-DSAD-Tau group exhibited enhanced proliferation, as indicated by a higher percentage of Ki67^+^/hN^+^ cells compared to the Cont-WT MG in the Cont-WT-DSAD-Tau group (Figure S6P, S6Q).

**Fig 6.**
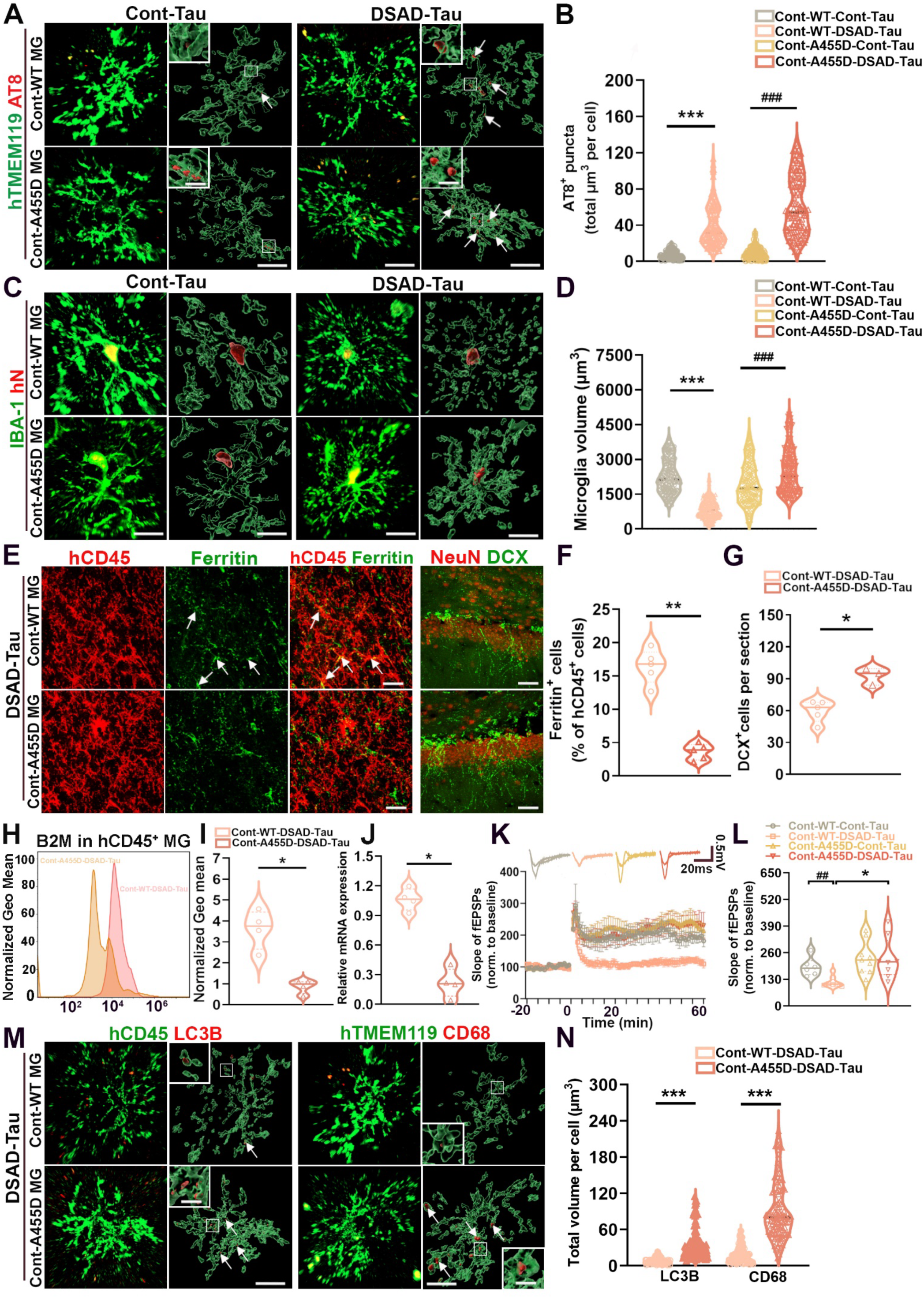
CSF2RB A455D mutation resilience is independent of trisomy 21. (A) Representative raw fluorescent super-resolution and 3D surface rendered images showing colocalization of hTMEM119+ and AT8+ staining in 6-month-old chimeric mice receiving injection of Cont or DSAD Tau at the age of 8 weeks. Scale bars: 5 μm and 1 μm in the original and enlarged images, respectively. (B) Quantification of AT8^+^ p-tau in Cont-WT and Cont-A455D MG following Cont and DSAD Tau injection (n = 125-128 from 3 mice per group). Two-way ANOVA test, \*\*\**P* < 0.001. Data are presented as mean ± SEM. (C) Representative images of IBA^+^/hN^+^ human microglia in Cont-Tau and DSAD-Tau groups. Scale bars:5 μm and 1 μm. (D) Quantification of microglia volumes in Cont-WT and Cont-A455D MG following Cont and DSAD Tau injection (n = 125-128 from 3 mice per group). Two-way ANOVA test, \*\*\**P* < 0.001. Data are presented as mean ± SEM. (E) Representative images showing colocalization of hCD45^+^ and Ferritin^+^ staining, DCX^+^ and NeuN^+^ in Cont-WT-DSAD-Tau and Cont-A455D-DSAD-Tau groups. Arrows indicate Ferritin^+^ and/or hCD45^+^ staining. Scale bar: 20 μm. (F) Quantification of the percentage of Ferritin in hCD45^+^ cells (n = 4-5). Student’s t-test, \*\**P* < 0.01. Data are presented as mean ± SEM. (G) Quantification of DCX^+^ cells (n = 3-4). Student’s t-test, \**P* < 0.05. Data are presented as mean ± SEM. (H) Flow cytometry analysis of B2M expression in Cont-WT-DSAD-Tau and Cont-A455D-DSAD-Tau groups. (I) Quantification of B2M expression in Cont-WT-DSAD-Tau and Cont-A455D-DSAD-Tau groups (n = 4). Student’s t-test, \**P* < 0.05. Data are presented as mean ± SEM. (J) qPCR analysis of B2M mRNA expression in chimeric mice at month 6 (n = 4 mice per group). Student’s t-test, \**P* < 0.05. Data are presented as mean ± SEM. (K) Representative traces of baseline and last 10 min fEPSP after 3X 100 Hz LTP induction. Quantification of LTP after LTP induction in 4 months old chimeras (n = 7-10 slices from 3-4 mice per group). (L) Quantification of the last 10 min of fEPSP slope after LTP induction in 4 months old chimeras (n = 6-9 slices from 3 mice per group). Two-way ANOVA test, \**P* < 0.05 and ^##^*P* < 0.01. Data are presented as mean ± SEM. (M) Representative images of hCD45 LC3B and hTMEM119 CD68 staining in 6-months-old Cont-A455D-DSAD-Tau and Cont-WT-DSAD-Tau chimeras. Arrows indicate LC3B^+^ and CD68^+^ puncta. Scale bars:5 μm and 2 μm in the original and enlarged images, respectively. (N) Quantification of LC3B^+^ and CD68^+^ puncta in hCD45^+^microglia (n = 121-123 microglia from 3 mice per group). Student’s t-test, \*\*\**P* < 0.001. Data are presented as mean ± SEM.

To further explore the protective function of CSF2RB A455D mutation in control microglia, we first compared the morphology of hN^+^ Cont-A455D and Cont-WT microglia following DSAD-Tau injection by double-staining IBA-1 and hN. The hN^+^/IBA-1^+^ Cont-A455D microglia in the DSAD-Tau group displayed increased microglia volume (Figure 6C, 6D). The hCD45 and ferritin staining showed that there was a decreased number of ferritin^+^/hCD45^+^microglia in the Cont-A455D-DSAD-Tau group, compared to Cont-WT-DSAD-Tau group (Figure 6E, 6F). Next, we analyzed the expression of B2M with flow cytometry and quantitative PCR and found increased expression of B2M at both protein and mRNA levels in Cont-WT-DSAD-Tau group, as compared to Cont-A455D-DSAD-Tau group (Figure 6H-6J). A significant increase of DCX^+^ newborn neurons in the hippocampus was also detected in Cont-A455D-DSAD-Tau group, compared to Cont-WT-DSAD-Tau group (Figure 6E, 6G). Furthermore, we assessed LTP induction in the hippocampal slices prepared from these chimeric mice. As shown in Figure 6K and 6L, Cont-WT-DSAD-Tau mice exhibited a significant reduction of fEPSP slope in comparison to Cont-WT-Cont-Tau mice, suggesting impaired hippocampal synaptic plasticity. In contrast, the enhancement of fEPSP slope persisted for 60 min in Cont-A455D-DSAD-Tau chimeras. All these LTP recording slices similarly showed a wide distribution of xenografted, hTMEM119^+^ human microglia (Figure S6R). As shown in Figure 6M, LC3B^+^ puncta were seen inside the hCD45^+^ Cont-WT-DSAD-Tau and Cont-A455D-DSAD-Tau microglia. Cont-A455D-DSAD-Tau microglia showed a higher volume of LC3B^+^ autophagosomes. We also found that Cont-A455D-DSAD-Tau microglia showed a higher volume of CD68^+^ phagolysosomes. Altogether, these *in vitro* and *in vivo* data illustrate that similar to its effect on DS microglia, CSF2RB A455D mutation consistently activates pSTAT5 in both control human iPSC-and ESC-derived microglia, upregulates phagocytosis and autophagy, and confers protection to A455D microglia and neuronal function against the detrimental effects induced by pathological tau.

### Control A455D microglia phagocytize and replace WT microglia in chimeric brains in response to pathological tau

To explore the interactions between A455D MG and WT MG within the same brain in response to pathological tau, we co-transplanted GFP^+^ CAGG-WT PMP and GFP-negative (GFP^-^) Cont-A455D PMP at a 1-to-1 ratio into the hippocampus of P0 immunodeficient mice. Subsequently, we administered Cont-Tau or DSAD-Tau injections at the age of 2 months (Figure 7A). After a 6-month cell engraftment period, both hN^+^/GFP^-^Cont-A455D and hN^+^/GFP^+^ CAGG-WT MG were broadly distributed throughout the hippocampus in both Cont-Tau and DSAD-Tau groups (Figure 7B). The majority of hN^+^ donor-derived cells (>90%) in both Cont-A455D+CAGG-WT-Cont-Tau and Cont-A455D+CAGG-WT-DSAD-Tau groups were hTMEM119^+^ (Figure 7C, 7F). Remarkably, by 6 months post-transplantation, there was a higher percentage of hTMEM119^+^/hN^+^/GFP^-^(Cont-A455D) MG compared to hTMEM119^+^/hN^+^/GFP^+^ (CAGG-WT) MG in the DSAD-Tau group, indicating A455D MG had outcompeted WT MG in the hippocampus in the setting of pathological tau challenge. In contrast, in the Cont-Tau group, there was no significant difference between the two populations of human microglia (Figure 7B,7C, 7G). In addition, the percentage of IBA-1^+^/hN^+^/GFP^-^(Cont-A455D) MG was higher than that of IBA-1^+^/hN^+^/GFP^+^ (CAGG-WT) MG in the DSAD-Tau group (Figure 7C, 7G), consistently suggesting that A455D MG had outcompeted WT MG in the hippocampus. To further assess the impact of DSAD-Tau on Cont-A455D and CAGG-WT MG in the same chimeric mouse brain, we compared the morphology of Cont-A455D MG and CAGG-WT MG following DSAD-Tau injection. As illustrated in Figure 7D and 7H, the hN^+^/IBA-1^+^/GFP^-^Cont-A455D MG displayed an increased cell volume in response to DSAD-Tau compared to hN^+^/IBA-1^+^/GFP^+^ CAGG-WT MG (Figure 7H). Additionally, we observed that Cont-A455D MG engulfed GFP^+^ debris in the DSAD-Tau group but not in the Cont-Tau group (Figure 7E, 7I). Moreover, Cont-A455D MG exhibited more LC3B^+^ puncta and a higher volume of CD68^+^ phagolysosomes than CAGG-WT MG in the DSAD-Tau group (Figure 7J-M).

**Fig 7.**
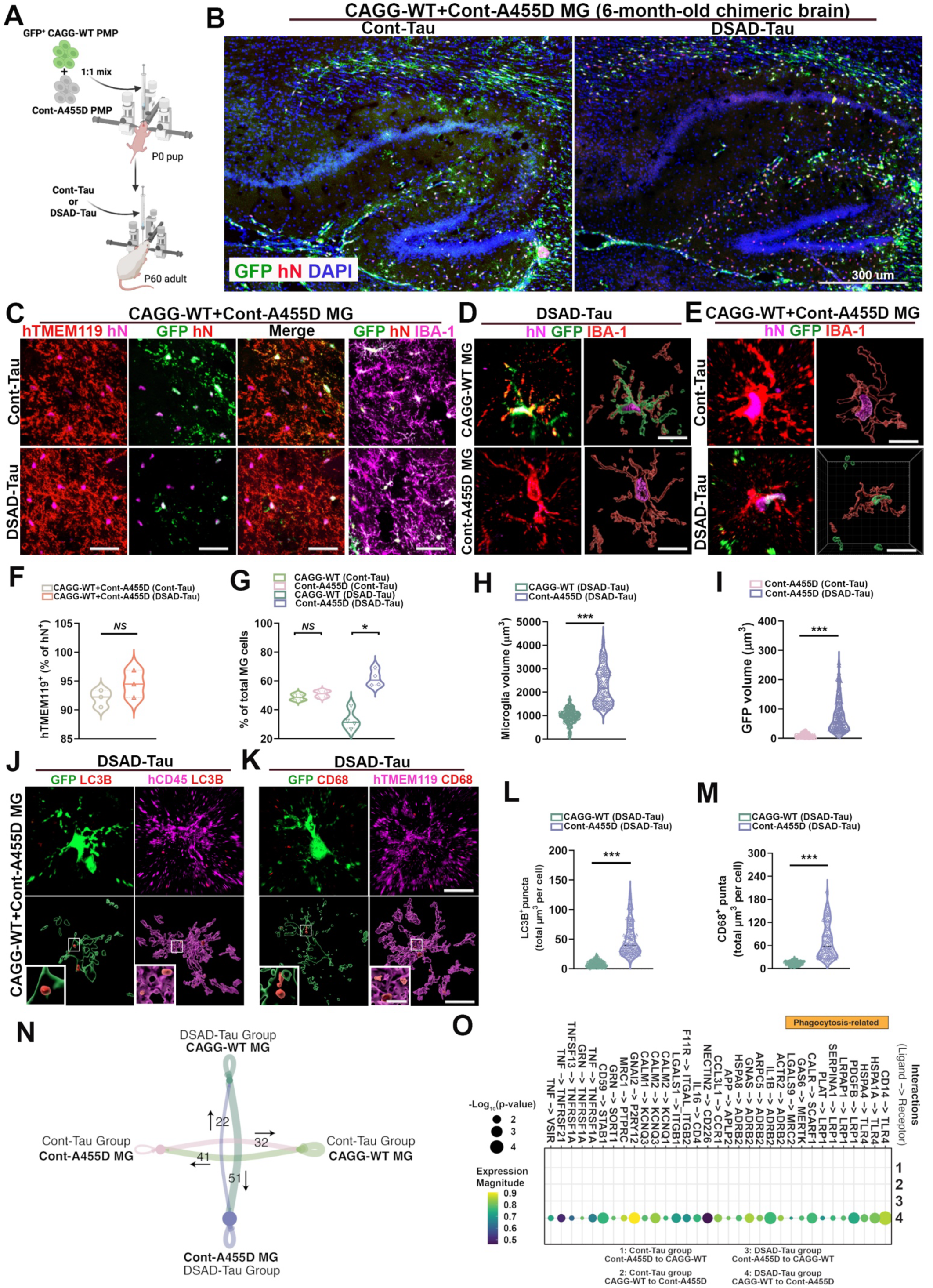
Control A455D microglia replace control WT microglia in response to pathological tau. (B) A schematic diagram showing the design of the co-transplantation experiment. (C) Representative images from sagittal brain sections showing the distribution of transplanted Cont-A455D and CAGG-WT at the age of 6-month-old chimeras. Scale bar: 300 μm. (D) Representative images showing colocalization of hTMEM119^+^ hN^+^, GFP^+^ hN^+^ and IBA-1^+^ hN^+^ staining in 6-month-old chimeras. Scale bar: 20 μm. (E) Representative images showing colocalization of IBA-1^+^ hN^+^ GFP^+^ and IBA-1^+^ hN^+^ GFP-staining in CAGG-WT-Cont-A455D-DSAD-Tau group. Scale bar: 5 μm. (F) Representative raw fluorescence super-resolution and 3D surface rendered images showing images showing colocalization of IBA-1^+^ hN^+^ and GFP^+^ staining in 6-month-old CAGG-WT-Cont-A455D chimeric mice receiving injection of Cont or DSAD Tau at the age of 8 weeks. Scale bar: 5 μm. (G) Quantification of the percentage of hTMEM119 in hN^+^ cells from 6-month-old chimeric mice (n = 3), Student’s t-test, *NS*, not significant. Data are presented as mean ± SEM. (H) Quantification of the percentage of CAGG-WT and Cont-A455D MG in total MG cells from 6-month-old chimeric mice (n = 3), Student’s t-test, \**P* < 0.05, *NS*, not significant. Data are presented as mean ± SEM. (I) Quantification of microglia volumes in CAGG-WT and Cont-A455D MG following DSAD-Tau injection (n = 132-135 from 3 mice per group). Student’s t-test, \*\*\**P* < 0.001. Data are presented as mean ± SEM. (J) Quantification of GFP^+^ volumes in Cont-A455D MG following Cont-Tau and DSAD-Tau injection (n = 125 from 3 mice per group). Student’s t-test, \*\*\**P* < 0.001. Data are presented as mean ± SEM. (K) Representative raw fluorescence super-resolution and 3D surface rendered images showing images of colocalization of LC3B^+^ in GFP^+^MG and hCD45^+^MG in 6-month-old CAGG-WT-Cont-A455D chimeric mice receiving injection of DSAD Tau at the age of 8 weeks. Scale bars: 5 μm and 2 μm in the original and enlarged images, respectively. (L) Representative raw fluorescence super-resolution and 3D surface rendered images showing images of colocalization of CD68^+^ in GFP^+^MG and hCD45^+^MG in 6-month-old CAGG-WT-Cont-A455D chimeric mice receiving injection of DSAD-Tau at the age of 8 weeks. Scale bars:5 μm and 2 μm in the original and enlarged images, respectively. (M) Quantification of LC3B^+^puncta in CAGG-WT and Cont-A455D MG following DSAD-Tau injection (n = 140 from 3 mice per group). Student’s t-test, \*\*\**P* < 0.001. Data are presented as mean ± SEM. (N) Quantification of CD68^+^puncta in CAGG-WT and Cont-A455D MG following DSAD-Tau injection (n = 125 from 3 mice per group). Student’s t-test, \*\*\**P* < 0.001. Data are presented as mean ± SEM. (O) Circle plot showing the number of inferred LR interactions between groups. Arrows indicate the direction of interactions (from ligands to receptors). (P) Dot plot showing the CAGG-WT-DSAD-Tau (ligands) to WT-A455D-Cont-Tau (receptors)-specific LR interaction pairs.

The co-transplantation of CAGG-WT and Cont-A455D MG offered an opportunity to investigate their interactions at molecular levels in response to the same brain environment. We then conducted scRNA-seq analysis on the chimeric mouse brains and performed cell-cell communication (CCC) analysis to explore ligand-receptor (L-R) interactions between CAGG-WT and Cont-A455D MG. After excluding mouse cells and implementing quality control measures, we found that in the DSAD-Tau group, a significant increase in L-R interactions was observed from CAGG-WT MG to Cont-A455D MG, a pattern not evident in the Cont-Tau group (Figure 7N). To specifically identify L-R interactions originating from CAGG-WT MG and targeting Cont-A455D MG in the DSAD-Tau group, we excluded common interactions seen in CAGG-WT MG to Cont-A455D MG in the Cont-Tau group, as well as Cont-A455D MG to CAGG-WT MG in the DSAD-Tau group. This analysis revealed 36 significantly distinct L-R pairs (Figure 7O). Importantly, receptors involved in microglial phagocytotic recognition of ‘eat-me’ signals, such as *MERTK, LRP1, SCARF1*, and *TLR4* ^66,67^, were uniquely inferred in interactions from CAGG-WT MG to Cont-A455D MG in the DSAD-Tau group, suggesting a unidirectional phagocytosis of CAGG-WT MG by Cont-A455D MG, specifically in the DSAD-Tau-treated group (Supplementary Table S11).

Furthermore, by comparing DEGs in DSAD-Tau groups, including CAGG-WT-DSAD-Tau vs. Cont-A455D-DSAD-Tau and DS-WT-DSAD-Tau vs. DS-A455D-DSAD-Tau, the Venn diagram showed 53 DEGs were commonly downregulated in the two datasets (Figure S7A). GO and Reactome enrichment analyses of the overlapped DEGs revealed that many were associated with type I and II interferon pathways (Figure S7B-S7C), similar to the observation in DS-A455D MG (Figure 4C). Comparing DEGs in DSAD-Tau groups, including CAGG-WT-DSAD-Tau vs. Cont-A455D-DSAD-Tau and DS-WT-DSAD-Tau vs. DS-A455D-DSAD-Tau, the Venn plot showed that 10.1% of DEGs were common, exclusively interferon-related genes. This indicates that in our DSAD-Tau model, CSF2RB A455D mutation in both DS MG and Cont MG similarly suppressed interferon signaling pathways after exposure to pathological tau (Figure S7A). Additionally, co-transplanting CAGG-WT and Cont-A455D MG revealed no difference in senescence gene enrichment in CAGG-WT+Cont-A455D-Cont-Tau and CAGG-WT+Cont-A455D-DSAD-Tau groups (Figure S7D). This suggests that the CSF2RB A455D mutation exerts protective functions in DSAD-Tau conditions. Taken together, these results demonstrate that Cont-A455D MG display upregulated phagocytosis and autophagy functions, engulfing and replacing CAGG-WT MG, providing protection against pathological tau-induced alterations.

## Discussion

Investigating mechanisms of resilience to AD may be key to identifying therapeutic interventions that have the potential to delay or halt the progression to AD dementia. However, this process has been challenging, in part due to the rarity of individuals who express such a phenotype. Recent milestone case report studies have identified two individuals from a family with a history of PSEN1 mutation-induced early-onset AD, suggesting that the APOE3 Christchurch (R136S) mutation and a variant in RELN (H3447R) may be responsible for resistance to autosomal dominant AD ^68,69^. In this study, utilizing hPSC-based *in vitro* microglia culture and *in vivo* human microglia chimeric mouse brain models, we present the following evidence demonstrating that a trisomy 21-associated CSF2RB A455D mutation specifically in microglia confers resilience to AD.

First, previous studies have demonstrated that soluble p-Tau extracted from human AD brain tissue triggers the microglial cell death *in vitro*^44^. We found that the CSF2RB A455D mutation provides protection to both DS and control hPSC-derived microglia against p-tau-induced cytotoxicity *in vitro* (Figure 1M, S6J). Secondly, the CSF2RB A455D mutation confers protection to DS and control microglia against senescence in the human microglia chimeric mouse brain, as indicated by improved morphology and decreased expression of the microglial senescence marker ferritin. scRNA-seq analysis of the chimeric mouse brain and experimental validation also demonstrated that DS and control microglia carrying the CSF2RB A455D mutation exhibit lower expression of senescence genes compared to DS and control microglia carrying the wildtype CSF2RB gene. Mechanistically, our previous study uncovered that exposure to pathological tau results in an upregulation of IFN-I signaling in DS-WT MG, playing a role in hastening the senescence of DS microglia^42^. As shown in our current scRNA-seq analysis, the CSF2RB A455D mutation suppresses the activation of IFN-I signaling, likely through the activation of JAK/STAT5, which is generally antagonistic to IFN-I receptor-activated STAT1 and STAT2 ^70^. This partly contributed to mitigated senescent and dystrophic changes in Cont-A455D and DS-A455D microglia. In addition, recent findings suggest that heightened autophagy prevents the senescence of microglial cells^61^. Consistent with this, we also observed the upregulated autophagy in DS-A455D MG compared to DS-WT MG, which further aids in preventing microglial senescence in response to pathological tau (Figure 4K-4P). Thirdly, the CSF2RB A455D mutation diminished the formation of microglia DAM state, characterized by the expression of genes associated with lipid droplet accumulation (Figure 3D). Furthermore, especially in DS microglia, the CSF2RB A455D mutation prompted the shift of homeostatic microglia to a more protective DAM microglial population, distinguished by the expression of genes related to tissue remodeling and repair (Figure 3C). Lastly, the CSF2RB A455D mutation in microglia protected neuronal functions against pathological tau, particularly hippocampal neurogenesis and synaptic plasticity, as evident by LTP recording (Figure 5F-5I). At the molecular level, our scRNA-seq in combination with experimental validation pinpoints B2M as a key molecule. Besides serving as a component of MHC class I molecules, B2M has been recognized as a secreted pro-aging factor predominantly expressed by microglia in the brain. This factor adversely affects synaptic functions, neurogenesis, and cognitive function ^62–64^. We demonstrate that the CSF2RB A455D mutation significantly decreases the expression of B2M in microglia induced by exposure to pathological tau (Figure 5A-5C). Moreover, the CSF2RB A455D mutation mitigated neuroinflammatory response and reduced the expression of inflammation-related genes in microglia. Furthermore, the neuroprotective effects conferred by A455D microglia are also attributed to their upregulated phagocytic function, as indicated by a higher volume of CD68+ phagolysosomes in A455D microglia than WT microglia, as well as enhanced autophagy functions in A455D microglia (Figure 4K-4P, 6M-6N). We found that A455D MG, in contrast to WT MG, exhibit more p62^+^ puncta and increased uptake of AT8^+^ p-Tau proteins (Figure 4M). Previous studies have demonstrated that the induction of p62, an autophagy receptor, is essential for the formation of α-synuclein/ubiquitin-positive puncta, which are subsequently degraded by autophagy^71^. Our results suggest that A455D MG likely possess enhanced phagocytosis and degradation of pathological tau, contributing to their neuroprotective effects.

The findings of this study carry significant implications for clinical translation and complement recent studies supporting the potential of microglia replacement therapy for treating AD^72–75^. It remains uncertain whether donor-derived cells, present in the diseased brain environment for an extended period, can sustain their protective function or undergo dysfunctional changes that cause detrimental effects ^75^. Indeed, concerns about the transfer of pathology from the host to the graft have been acknowledged in the context of developing cell replacement therapy for Parkinson’s disease, where previous studies have documented cytoplasmic deposits of phosphorylated α-synuclein in a subset of grafted dopaminergic neurons in both animal models and patients^76–78^. Given the high sensitivity of microglia to their environment, it is crucial to identify genes that confer resilience, enabling microglia to withstand AD pathology while preserving their normal functions and providing enduring protection to neurons ^1^. Our results reveal that the CSF2RB A455D functions as such a gene variant in human microglia.

Acting as a component of the GM-CSF receptor, the beneficial effects of the CSF2RB A455D mutation and the resultant activation of CSF2R-mediated signaling in microglia are consistent with preclinical and clinical studies^32–36^ indicating that GM-CSF treatment stimulates the innate immune system and enhances cognition. However, somatic mutations in HSCs can induce clonal hematopoiesis, elevating the risk of blood cancer and other diseases, such as cardiovascular diseases^79,80^. Similarly, gain-of-function mutations in CSF2RB, such as the A445D mutation, represent clonal variants responsible for ML in a subset of DS individuals ^29^. Fortunately, microglia have never been identified as glioma-initiating cells, and in this study, we did not observe any tumorigenicity of A455D microglia, as evidenced by a similar proliferation rate between WT and A455D microglia in chimeric mice receiving Cont-Tau. While A455D microglia exhibited a higher proliferation rate than WT microglia in chimeric mice receiving DSAD-Tau, we did not observe any cell overgrowth or tumor formation in the chimeric brain for over 6 months of age. Therefore, delivering A455D microglia to the brain may provide long-lasting beneficial effects and mitigate the risk of inflammation associated with the systemic administration of GM-CSF^81^ and the potential tumorigenicity if A455D hematopoietic progenitors are peripherally transplanted.

Using human macroglial chimeric mouse brain models, a recent study demonstrated that aged and dysfunctional human macroglia in diseased brains could be effectively eliminated and replaced by intracerebral delivery of young allogeneic human macroglial progenitors^82^. Different from macroglia, microglia are professional phagocytes with a primary role in eliminating dysfunctional, diseased, or apoptotic cells in the brain. We observe that when co-transplanted into the same brain, A455D microglia engulf WT microglia in response to pathological tau, outcompeting and replacing the WT microglia in the chimeric brains. Although A455D microglia and WT microglia were generated from hiPSC and hESC, respectively, the replacement was unlikely due to a growth advantage of the hPSCs because the A455D MG and WT MG comprised a comparable proportion of total hN+ donor-derived cells in the absence of pathogenic tau. The engulfment of WT microglia by A455D microglia following exposure to p-Tau is additionally substantiated by scRNA-seq ligand-receptor interaction analysis of the interplay between WT and A455D microglia. As demonstrated in recent studies^72,73,83,84^, efficient microglia replacement by peripheral cell sources or through direct microglia transplantation requires the treatment with destructive agents, such as CSF1R inhibitors and/or toxic preconditioning of the bone marrow niche. Our results suggest that in the context of AD, eliminating endogenous senescent/dystrophic microglia may not be necessary. This is also anticipated in people with clonal hematopoiesis and resilience to AD, given its nature of somatic mutations instead of germline mutations. In such individuals, myeloid cells originating peripherally with acquired protective mutations are likely to enter the brain, where they differentiate into microglia-like cells and outcompete the endogenous senescent/dystrophic microglia^6,7,85^. Hence, our present results indicate the potential application of hiPSC-derived CSF2RB A455D microglia for developing efficient autologous or allogeneic microglial replacement therapy through direct brain transplantation, aimed at treating AD and other age-related neurodegenerative disorders.

The exceedingly rare detection of DS individuals exhibiting AD resilience in clinical settings, coupled with the limited accessibility of functional brain tissue from such DS individuals, hinder the isolation of myeloid cells, potentially from brain regions with heightened blood-brain barrier permeability, a process currently essential for rare-variant analysis ^6^. While this presents substantial challenges to establish a correlation between CSF2RB A455D and AD resilience in patients, we demonstrate in human-mouse chimeric brain model that engrafted A455D microglia eliminate and outcompete the senescent/dystrophic WT microglia and provide protection to neuronal function. Previous studies from us and others demonstrate that human-mouse chimeric brain models recapitulate the degenerative phenotypes of human neurons and glia observed in human AD brain tissue^37,41,42,86^. These results pave the way for utilizing the chimeric model in the current study to assess the protective effects of CSF2RB A455D microglia against effects driven by pathological tau, which is much more closely related to cognitive decline as compared to Aβ pathology. In future studies, creating immunodeficient AD mouse models of tauopathy to receive human CSF2RB A455D microglia transplantation can aid in further examining the responses of A455D microglia in brain environments with more widespread tau pathology.

## Acknowledgements

This work was in part supported by grants from the NIH (R01NS102382, R01NS122108, and R01AG073779 to P.J.). M.J. was supported by a postdoctoral fellowship award from the New Jersey Department of Health (CAUT24DFP004). We thank the UCI-ADRC, which is funded by NIH/NIA Grant P30AG066519 and the Brightfocus Foundation (BFF17-0008), for providing us with DSAD and control human brain tissues. Y.L. was supported by NIH R01NS110707. Additional support came from the NIH (R01AG064579 and RF1NS128800), the JSRM Foundation and the Alzheimer’s Association to S.F. We thank Dr. Ping Xie from Rutgers University for her suggestions on p-STAT5 flow cytometry experiments. We appreciate Dr. Kelvin Kwan from Rutgers University for aiding in library preparation for scRNA-seq. We are also thankful to Ava Papetti, Anee Kumar, and Kushal Aluru from the Jiang laboratory for their assistance with immunohistochemistry. We thank Françoise Chanut for editorial assistance.

## Author Contributions

M.J. and P.J. designed experiments and interpreted data; M.J. carried out most of experiments with technical assistance from H.Z. and R.D.; Z.M. performed RNA-seq data analyses and assisted with sequencing data interpretation; R.D. performed electrophysiological recordings; J.P. and E.H. prepared human brain tissue extracts; Y.L. and H.X. generated the CSF2RB A455D DS and CAGG hPSC lines; S.F. and E.H. provided critical suggestions to the study; P.J. conceived the concepts, directed the project, and wrote the manuscript together with M.J. and input from all co-authors.

## Methods

### Construction of CSF2RB WT and A455D mutant targeting vectors

Human CSF2RB WT (exons 11 to 14) and A455D mutant donor targeting sequences were cloned into pDONR221 (ThermoFisher) with the following layout (Supplementary Figure S1): 5’ homology arm - splice acceptor (SA) - cDNA fragment of exon 11 through exon 14 of WT or CSF2RB A455D mut - human growth hormone (hGH) poly(A) signal - floxed EGFP (or mRuby) - P2A - puromycin resistance cassette driven by the EF1α promoter - 3’ homology arm. The donor vectors were designed to integrate to intron 10 of the CSF2RB genomic locus. The 5’ and 3’ HAs were amplified using hPSC WA09 genomic DNA as a template. Sequences of corresponding PCR primers (CSF2RB 5’HA-F, CSF2RB 5’HA-R, CSF2RB 3’HA-F, and CSF2RB 3’HA-R) for amplifying HAs are listed in Supplementary Table S1). The 5’ and 3’ homology arms (HA) were 708 bp and 634 bp in length, respectively. To generate the codon optimized cDNA fragments (exons no. 11-14) of human CSF2RB WT and A455D, Addgene vectors (cat. no. 125750 and cat. no. 125751) were used as templates for PCR amplification (PCR primers SA-CSF2RB-Ex11-14-F and CSF2RB-Ex11-14-R are listed in Supplementary Table S1).

As shown in the plasmid layout (Supplementary Figure S1), the portion of the codon optimized WT CSF2RB cDNA fragment that was comprised of exons 11-14 was preceded by a synthetic splice acceptor (SA) sequence and followed by an hGH polyA sequence. For the CSF2RB A455D mutant donor, the codon optimized cDNA sequences contained GCT-GCC mutations which encodes aspartic acid (D) instead of alanine (A) for amino acid no. 455. Donor sequences of both WT and mut were mammalian codon optimized which have multiple silent mutations that reduce the possibility of premature cross-over events during homology directed repair (HDR) during CRISPR-Cas9 editing. P2A is the self-cleaving peptide that allows for simultaneous, separate protein expression of EGFP (or mRuby) cassette and puromycin resistance fragment. All fragments were assembled by Gibson assembly (New England Biolabs) as described previously ^87,88^ The final constructs were selected with kanamycin and named H598 (WT GFP), H599 (A455D GFP), H628 (WT mRuby), and H629 (A455D mRuby). Note that the purpose of including a floxed EGFP or mRuby expression cassette, connected by P2A with a puromycin-resistance fragment, was to increase the efficiency of identifying correctly targeted clones in hPSCs. Upon further differentiation, expression of EGFP and mRuby was completely silenced and undetectable.

### Construction of single guide RNA vector for the CRISPR/Cas9 system to target the CSF2RB locus

The pX330-U6-Chimeric_BB-CBh-hSpCas9 vector ^89^, which allows for human single guide RNA (sgRNA) expression together with mammalian codon-optimized *Streptococcus pyogenes* Cas9 expression, was a gift from Feng Zhang (Addgene plasmid # 42230; http://n2t.net/addgene:42230; RRID:Addgene_42230). The sequence for making sgRNA for mediating CSF2RB targeting was located within the intron 10 of CSF2RB: 5’ CAGAACAAGACAAGTCCAAG**GGG** 3’ (PAM sequence is in bold). To construct the sgRNA expression vector, a pair of oligos (CSF2RB-f3-top 5’ CACCCAGAACAAGACAAGTCCAAG 3’ and CSF2RB-f3-bottom, 5’ AAACCTTGGACTTGTCTTGTTCTG 3’) was designed using Benchling, a cloud-based biology software (https://www.benchling.com). The oligos were synthesized and subcloned into pX330 sgRNA-Cas9 2-in1-expression vector following a previously published protocol ^90^.

### T7 Endonuclease I (T7EI) assay

T7EI assay was performed as previously described ^91–93^. Briefly, 1.5×10^5^ 293FT cells (one well of a 12-well plate) were transfected with 1 µg CSF2RB -sgRNA expression vectors and 1 µg Cas9 expression vector, or GFP (control) vector, using Lipofectamine 3000 (Life Technologies). Cells were harvested 72 h post transfection, digested with 100 µL genomic DNA quick extraction solution (Fisher), and incubated at 68°C for 15 min, and then 95°C for 8 min. The concentration of the genomic DNA was adjusted to 200 ng/µL before it was PCR amplified using Herculase II (Agilent). Approximately 400 ng purified PCR product (PCR purification kit, Qiagen) was denatured by incubating at 95°C for 5 min and the temperature was slowly lowered to < 30°C to allow the formation of heteroduplex. T7EI assay reaction was setup as follows: 400 ng hybridized DNA, 2 µL Digestion Buffer 2, and 1 μL (10 units/μL) T7EI enzyme. Reaction was incubated at 37°C for 20 min. Digested product was separated by 2.5% agarose gel. Gel bands were quantified using Image J and the cleavage efficiency (percentage of insertion and deletion, abbreviated as indel) was calculated following a previously reported protocol ^90,94^.

### Generation of the CSF2RB WT or A455D Mut hPSC using DS iPSC or control CAGG3.1 lines

The CSF2RB A455D induces STAT5 phosphorylation in a dominant-negative manner, as this mutation promotes dimerization of the transmembrane domains, either via homomeric interaction with another CSF2RB chain or heteromeric interaction with partner α chains of the IL-3, IL-5, and GM-CSF receptors. This results in an active conformation for downstream JAK signaling and STAT5 phosphorylation^29^. Hence, in this study, we generated hiPSCs homozygous for the mutation. Human PSC lines were maintained in chemically defined mTeSR plus medium (Stemcell Technologies Inc.) in a feeder free fashion and passaged every 4∼5 days at a 1:4∼1:6 ratio using ReLeSR (Stemcell technologies) as described previously ^88,95^. Alternatively, cells were passaged every 4∼5 days with Accutase (Innovative Cell Technologies) onto Matrigel (BD Biosciences)-coated culture plates at a ratio of 1:4∼1:8 supplemented with ROCK inhibitor Y-27632 (10 mM, R&D systems). To monitor and ensure the genetic stability of the cells, routine karyotyping examination was done every 10 to 15 passages. To generate CSF2RB WT or A455D Mut hPSC lines hPSCs CAGG3.1 ^96^ or Down syndrome patient derived iPSCs DS2 ^42,97,98^, approximately 1×10^5^ hPSCs were plated on a well of a 12-well plate in mTeSR plus medium, one day prior to transfection. Cells from each well were co-transfected with CSF2RB (WT or A455D, 0.5 µg) and sgRNA-Cas9 (0.5 µg) vectors using Lipofectamine Stem (Life Technologies) following the manufacturer’s instructions. ROCK inhibitor (10 μM) was added for the first 24 h post-transfection to enhance single cell survival. Cells were fed with fresh mTeSR plus medium every day until Day 3 post transfection, when Puromycin (0.75-1 µg/mL) selection was started. Resistant single clones were isolated and manually picked after two to four days of selection and subsequently expanded individually. Genomic DNA extracted from single clones were examined by PCR for identification of correctly targeted hPSC clones. Primer sequences for CSF2RB-test 5-F and CSF2RB-test 5-R are listed in Supplementary Table S1. The PCR products with the correct size were verified by Sanger sequencing.

### Generation of the CSF2RB A455D Mut in control hiPSCs

sgRNAs were designed, synthesized, and chemically modified by Synthego. Donors with homology arms were designed and synthesized. Arms were designed to have sufficient length to ensure high knock-in efficiency. Silent mutations were included for each guide to reduce cutting post-editing and maximize the efficiency of the knock-in. Donor templates were provided either as single-stranded oligodeoxynucleotides (ssODNs) or plasmids. sgRNA sequence, test primers and donor sequence are shown in Supplementary Table S2. Control ND2.0 iPSCs were maintained in mTeSR plus as described above. CSF2RB-specific guide RNAs were complexed together with the Streptococcus pyogenes (SpCas9) to form a ribonucleoprotein (RNP). RNPs and donors were then delivered to the ND2.0 iPSCs via electroporation. Resulting clones are verified using Sanger sequencing.

### Summary of hPSC A455D mut lines used in the current study

Three parental hPSC lines were used in this study (Supplementary Table S3): (1) ND2.0 is an hiPSC line with normal karyotype derived from a healthy individual. The cells were originally obtained from Center for Regenerative Medicine, National Institutes of Health (https://commonfund.nih.gov/stemcells/lines). (2) Trisomy 21 DS2 line is an hiPSC line reprogrammed from DS patient fibroblasts (Coriell Institute for Medical Research). The iPSC reprogramming was performed in house. The DS2 line, together with other normal or DS iPSCs, has been described and characterized previously ^97^. (3) CAGG3.1 ^96^ is an hESC GFP reporter cell line, with a CAG promoter-driven eGFP cassette integrated at the chromosome 13 safe harbor locus of the WA09 hESCs, which offers a convenient GFP reporter for direct visualization when the hESCs are differentiated into terminal cell types (e.g., microglia in the current study) or grafted in vivo.

### hiPSC lines generation, culture, and quality control

A total of six hiPSC lines were used in this study: one pair of DS-WT and DS-A455D, and two pairs of Cont-WT and Cont-A455D (Supplementary Table S3). The hiPSC lines were fully characterized and completely de-identified ^97,98^. All hiPSCs were cultured on dishes coated with hESC-qualified Matrigel (Corning) in mTeSR plus media (STEMCELL Technologies) under a feeder-free condition. The hiPSCs were passaged with ReLeSR media (STEMCELL Technologies) once per week.

### Differentiation and culture of PMPs

PMPs were generated from the three pairs of Cont and DS hiPSC cell lines using a previously established protocol ^99^. The yolk sac embryoid bodies (YS-EBs) were generated by treating the YS-EBs with mTeSR 1 media (STEMCELL Technologies) supplemented with bone morphogenetic protein 4 (BMP4, 50 ng/ml), vascular endothelial growth factor (VEGF, 50 ng/ml), and stem cell factor (SCF, 20 ng/ml) for 6 days. To stimulate myeloid differentiation, the YS-EBs were plated on dishes with X-VIVO 15 medium (Lonza) supplemented with interleukin-3 (IL-3, 25 ng/ml) and macrophage colony-stimulating factor (M-CSF, 100 ng/ml). At 4-6 weeks after plating, human PMPs emerged into the supernatant and were continuously produced for more than 3 months.

### *In vitro* differentiation of PMPs to microglia

PMPs were differentiated in the medium composed of DMEM/ F12 supplemented with N2, 2 mM Glutamax, 100 U/mL penicillin and 100 mg/mL streptomycin, 100 ng/mL M-CSF (Peprotech), 100 ng/mL IL-34 (Peprotech), and 10 ng/mL GM-CSF (Peprotech) for two weeks ^99^. The medium was changed once a week.

### Animals and cell transplantation

All animal work was performed without sex bias with the approval of the Rutgers University Institutional Animal Care and Use Committee. PMPs were collected from the supernatant and suspended at a concentration of 100,000 cells/µl in PBS. These PMPs were subsequently injected into the brains of P0 Rag2^−/−^hCSF1 immunodeficient mice (C;129S4-*Rag2^tm^*^1^*^.1Flv^ Csf1tm1^(CSF1)Flv^ Il2rg^tm1.1Flv^/J*, The Jackson Laboratory) as previously described ^42,49,100^. The transplantation sites were bilateral from the midline = ±1.0 mm, posterior from bregma = −2.0 mm, and dorsoventral depths = −1.5 and −1.2 mm ^48^. All pups were placed in ice for 4-5 mins to anesthetize. The pups were then injected with 0.5 μl of cells into each site (four sites total), using a digital stereotaxic device (David KOPF Instruments) that was equipped with a neonatal mouse adapter (Stoelting). The pups were weaned at three weeks and kept for further experimentation at various time points.

### Preparation of soluble S1 fractions

The soluble S1 fractions from human samples (provided by the University of California Alzheimer’s Disease Research Center (UCI-ADRC) and the Institute for Memory Impairments and Neurological Disorders) were prepared following established procedures^44^ (Supplementary Table S4). Human tissues were homogenized in TBS (20 mM Tris-HCl, 140 mM NaCl, pH 7.5) containing protease and phosphatase inhibitors (Roche). Following homogenization, the samples were ultracentrifuged (4 °C for 60 min) at 100,000×*g* (Optima MAX Preparative Ultracentrifuge, Beckman Coulter). Supernatants, S1 fractions, were aliquoted and stored at −80 °C.

### Tau quantification by ELISA

The quantification of the total amount of soluble tau in S1 fractions was performed by using an ELISA kit (human Tau, Invitrogen), following the manufacturer’s protocol. The ELISA experiments were conducted in three independent experiments using triplicate replicas, and the concentration of each sample was listed in Supplementary Table S5.

### Intracerebral adult brain injection

All adult brain injections were performed using a Kopf stereotaxic apparatus (David Kopf, Tujunga, CA). Stereotaxic surgery was performed on two months old transplanted Rag2^−/−^hCSF1 immunodeficient mice. The mice were aseptically injected with human brain extracts (Cont-Tau or DSAD-Tau) in the dorsal hippocampus and the overlying cortex (bregma: −2.5 mm; lateral: +2 mm; depth: −2.4 mm and −1.8 mm from the skull). A dose of tau at 3.2 µg, which was previously shown to induce tau pathology in tau transgenic mice ^101–103^, was injected into each human microglial chimeric mouse. Concentrations of tau per injection site were 0.8 μg/µl each site for both DSAD-Tau and Cont-Tau. The mice were injected with 1 μl of Cont or DSAD-Tau into each site (four sites total). After the injection, the mice were kept for further experiments.

### Flow cytometry

Single-cell suspensions from the chimeric brains were washed and suspended in PBS with 1% BSA. The cells were blocked with rat serum and FcR blocking Ab (2.4G2), and then incubated with Ab conjugated to different fluorochromes for surface staining. Intracellular markers were stained after fixation and permeabilization of cells using an Intracellular or Phospho-Flow Staining kit (BD Biosciences). FACS data were acquired on a Northern Lights spectral flow cytometer (Cytek, Fremont, CA). The results were analyzed using the FlowJo software (TreeStar, San Carlos, CA)

### RNA isolation and quantitative reverse transcription PCR

Total RNA was extracted using TRIzol reagent (Thermo Fisher Scientific, 15596026), and RNA was reverse transcribed into complementary DNA (cDNA) using TaqMan™ Reverse Transcription Reagents (Thermo Fisher Scientific; N8080234). Total DNA was prepared with Superscript III First-Strand kit (Invitrogen). Real-time PCR was performed on the ABI 7500 Real-Time PCR System using the TaqMan Fast Advanced Master Mix (Thermo Fisher Scientific). All primers are listed in Table S12. The 2^−ΔΔ*Ct*^ method was used to calculate relative gene expression after normalization to the *GAPDH* internal control.

### Extracellular hippocampal slice recording

All the mice used in field recordings were 4 to 6-month-old and conducted at the Schaffer collateral-commissural pathway. The detailed procedures for slices recording were described previously ^42^. In brief, the mouse brains were quickly separated and put into ice-cold cutting solution (in mM): 206 Sucrose, 11 D-Glucose, 2.5 KCl, 1 NaH_2_PO_4_, 10 MgCl_2_, 2 CaCl_2_, and 26 NaHCO_3_ saturated with 95% O_2_/5% CO_2_, then cut into 360 μm slices perpendicular to the long axis of the hippocampus with Leica Vibratome (Leica, VT1200). The slices were recovered in recording solution (in mM): 120.0 NaCl, 3.0 KCl, 1.2 MgSO_4_, 1.0 NaH_2_PO_4_, 26.0 NaHCO_3_, 2.0 CaCl_2_, 11.0 D-glucose saturated with 95% O_2_/5% CO_2_ at 33°C for 30 min, and then transferred to room temperature for at least 1 hour before recording. The slices were recorded in a chamber perfused with recording solution saturated with 95% O_2_/5% CO_2_. For LTP recordings, synaptic responses were evoked by stimulation at 0.067 Hz at Schaffer collaterals and recorded with glass pipettes (3-4 MΩ) filled with recording solutions at stratum radiatum of CA1 region. LTP were induced by three trains of High-frequency stimulation (HFS, 100Hz, 1s) with 10 s intertrain interval. The last 10 min were used for analysis compared with baseline. All data were collected and analyzed with pCLAMP 10.7 (Molecular Devices), n represents the number of slices and N represents the number of animals. Usually, two to four slices per animal were used.

### Cell viability test

Cell viability was measured using the cell counting kit-8 (CCK-8) (Sigma-Aldrich, St. Louis, MO, USA) as previously described ^104^. In brief, the differentiated MG were seeded into a 96-well microplate and then were incubated with different treatments. Subsequently, 10 µL of CCK-8 solution was added to 100 µL media per well and incubated at 37 °C for 3-4 h. The colorimetric assay was measured using a SpectraMax iD3 microplate reader (Molecular Devices, San Jose, CA, USA).

### Tissue immunostaining, image acquisition, and analysis

Mouse brains were fixed with 4% paraformaldehyde. Mouse brains were placed in 20% and later in 30% sucrose for dehydration. Following dehydration, brain tissues were immersed in OCT and frozen for sectioning. The frozen tissues were cryo-sectioned with 30-μm thickness for immunofluorescence staining. The tissues were blocked with a blocking solution (5% goat or donkey serum in PBS with 0.8% Triton X-100) at room temperature (RT) for 1 hour. The primary antibodies were diluted in the same blocking solution and incubated with the tissues at 4 °C overnight (all the primary antibodies are listed in Supplementary Table 13). The sections were washed with PBS and incubated with secondary antibodies for 1 hour at RT. After washing with PBS, the slides were mounted with anti-fade Fluoromount-G medium containing 1, 40,6-diamidino-2-phenylindole dihydrochloride (DAPI) (Southern Biotechnology). Mouse brains older than four months underwent TrueBlack treatment following immunostaining, as detailed in our previous report using 0.4X TrueBlack treatment solution (Biotium)^100^.

All images were captured with a Zeiss 800 confocal microscope. Large scale images in Fig. 2A and 7B were obtained by confocal tile scan by the Zen software (Zeiss). To obtain a 3D reconstruction, images were processed by the Zen software (Zeiss). To visualize phagocytic function, super-resolution images in Figures. 2, 4, and 6-7 were acquired by Zeiss Airyscan super-resolution microscope at 63X with 0.2mm z-steps. To generate 3D-surface rendered images, super-resolution images were processed by Imaris software (Bitplane 9.9). To visualize the engulfment of AT8^+^ within microglia, CD68^+^ phagolysosome inside microglia, p62 and LC3B fluorescence found outside of the microglia were subtracted from the image via the mask function in Imaris software. The number of positive cells from each section was counted after a Z projection. The dystrophic (Iba-1^+^hN^+^) microglial morphological changes after tau injection were assessed by calculating the soma size and process length using the software Fiji with MorpholibJ plugin and integrated library as described before ^46,105^. Soma size was measured with a closing morphological filter connecting dark pixels and process length was analyzed with Skeletonize function.

### Bulk RNA-sequencing

PMPs generated from the three pairs of Cont and DS hiPSC lines were used for RNA extraction and RNA sequencing sample preparation. Total RNA was prepared with an RNAeasy kit (QIAGEN) (Chen et al., 2014) and libraries were constructed by using 600 ng of total RNA from each sample and utilizing a TruSeqV2 kit from Illumina (Illumina, San Diego, CA) following the manufacturer’s suggested protocol. The libraries were subjected to 75 bp paired read sequencing using a NextSeq500 Illumina sequencer to generate approximately 30 to 35 million paired-reads per sample. Sequenced reads were QC’ed by FastQC (v0.12.1) and trimmed for adaptor sequence/low-quality sequence using Trim Galore (version 0.6.10). Trimmed sequence reads were mapped to GRCh38 reference genome and GENCODE v44 primary assembly using STAR (version 2.7.11a). Read count extraction and normalization were performed using RSEM (version 1.3.1). DEG analysis was performed using DESeq2 R package (version 1.42.0). Linear regression and correlation analysis were performed using the arithmetic mean of variance stabilizing transformation (VST) normalized counts among all replicates of each experimental group.

### Single-cell RNA-sequencing

Chimeric mice were anesthetized and perfused with cold DPBS (1x). Brains were dissected and dissociated using Adult Brain Dissociation Kit (Miltenyi) according to the manufacturer’s instructions. Human cells were isolated by mouse cell depletion kit (Miltenyi) following the manufacturer’s protocols. Isolated cell suspension then loaded to the Chromium controller for scRNA-seq library preparation, using Chromium Next GEM Single Cell 3’ Kit v3.1 and Dual Index Kit TT Set A, following the manufacturer’s protocols. Quality control of the scRNA-seq libraries were performed by Agilent 2100 Bioanalyzer, and libraries were sequenced in Novaseq 6000 S4 flow cells.

### scRNA-seq data pre-processing

Raw reads were mapped by human (GRCh38, Ensembl 98, GENCODE v32) and mouse (mm10, Ensembl 98, GENCODE vM23) reference genome by Cell Ranger (7.1.0), with EGFP gene built in if needed. Cells were categorized as human cells if >75% genes were mapped to the human reference genome. Filtered human cells were kept for further analysis with gene number between 200 and 2500, and mitochondria content less than 10%. Downstream analysis was performed by Seurat R package (v4.3.1). Data was log-normalized and scaled with top 2,000 variable features. Linear (PCA) and non-linear (UMAP) dimensional reductions were performed using top 15 principal components (PCs). Graph-based clustering was performed using resolution = 0.3. Differential gene expression analysis was performed by the *FindMarkers* function using a negative binomial generalized linear model.

### Cluster similarity analysis

Cosine similarity scores were calculated to measure the transcriptomic similarity between clusters using the *CellCorHeatmap()* function provided by the SCP R package. Jaccard similarity index was used for cross-sample cluster comparisons. Cluster-specific marker genes were computed by Seurat *FindAllMarkers()* function, and filtered with avg_log2FC > 0.25 and adj.pvalue < 0.05. Jaccard similarity coefficients were calculated using the cluster-specific marker genes provided by Sun et al.^54^, and this study by GeneOverlap R package.

### Enrichment analysis and Gene Set Enrichment Analysis (GSEA)

Enrichment analysis was performed using ClusterProfiler R package. Differentially expressed genes with abs(avg_log2FC) > 0.25 and adj.pvalue < 0.05 were used for gene ontology (GO), Kyoto Encyclopedia of Genes and Genomes (KEGG), and Reactome pathways enrichment analysis. Over Representation Analysis (ORA) was performed to enrich provided gene lists, using all other genes in the database as background. P-values were calculated by hypergeometric distribution. Log2FC-ranked gene lists were used for GSEA input, and GSEA was performed by the *fgsea* algorithm.

### Pseudotime trajectory inference

Pseudotime analysis was performed using the *slingshot* algorithm wrapped in the SCP R package. Analysis was performed separately using cells from different experimental groups by RunSlingshot() function. To visualize gene expression changes across inferred pseudotime, heatmaps showing dynamic features were plotted by DynamicHeatmap() function, dynamic features across pseudotime were plotted using DynamicPlot() function.

### Cell-cell communication analysis

Cell-cell communication (CCC) analysis was performed using LIANA, by combining different ligand-receptor interaction prediction algorithms and CCC databases to make robust CCC predictions. liana_wrap() function was executed to perform CCC prediction with default parameters. Predicted CCC results were aggregated by the robust rank aggregation (RRA) method, provided by the liana_rank() function.

### Statistical analysis

All data are represented as mean ± SEM. When only two independent groups were compared, significance was determined by using two-tailed unpaired t-test with Welch’s correction. When three or more groups were compared, two-way ANOVA test was used. A p-value of < 0.05 was considered significant. All the analyses were done in GraphPad Prism v.9. All experiments were independently performed at least three times with similar results.

